# Expectation of aversive outcomes drives multimodal integration of behavioural, physiological and neural activations during human fear retrieval and generalisation

**DOI:** 10.64898/2026.01.09.698570

**Authors:** Leonie Rumpf, Madeleine Müller, Jan Haaker

## Abstract

Classical fear conditioning describes how neutral cues acquire a threat value, yet how learned associations are retrieved and generalised across similar stimuli specifically is an ongoing debate. We combined behavioural ratings, physiological measures, and fMRI in a two-day classical fear conditioning paradigm to characterize acquisition, retrieval, and generalisation across modalities. Twenty-five healthy participants completed acquisition trials on Day 1 and retrieval and generalisation trials on Day 2 using conditioned (CS+, CS-) and graded generalisation stimuli (GS). Outcomes included trial-wise US expectancy ratings, pre/post fear and arousal ratings, skin conductance responses (SCR), pupil dilation, and ROI-based fMRI (amygdala, hippocampus, insula, periaqueductal gray (PAG), locus coeruleus). Acquisition yielded robust CS+/CS-discrimination in behavioural ratings and increased BOLD responses in bilateral insula and PAG. During retrieval, US-expectancy ratings indicated early retrieval of CS contingency. The fMRI results showed greater BOLD activity during CS+ presentations than during CS-presentations in bilateral hippocampus, left insula and right PAG. Additionally, hippocampus-insula coupling increased. Critically, parametric modulation during retrieval revealed that trial-wise mean US-expectancy modulated BOLD responses in left insula and right PAG, with a trend in left hippocampus. Across generalisation, US-expectancy and pupil dilation responses followed graded profiles, which could be explained by a Gaussian model, whereas SCR generalised, but was not captured by a Gaussian model. Parametric modulation by US-expectancy correlated with BOLD activity in left PAG, with a trend in right hippocampus. Stimulus identity explained variance in bilateral insula and left PAG. Findings converge on a hippocampus-insula-PAG network that retrieves learned predictions, and scales defensive output according to similarity-based threat probability, linking subjective, physiological, and neural outcomes.

## INTRODUCTION

Threat learning paradigms, like classical fear conditioning, allow examining neurobiological responses to investigate how organisms learn to predict and respond to threats. It entails learning to predict the occurrence of an aversive unconditioned stimulus (US) by the presence of a preceding neutral cue (conditioned stimulus, CS+). The resulting expectation of the US during CS presentations elicits conditioned responses across behavioural, physiological, and neural outcome measures. In humans, this process of acquisition engages a distributed cortico-limbic-brainstem network including the amygdala, hippocampus, insula and periaqueductal gray (PAG) (Mobbs et al., 2009; Phelps et al., 2001; Sehlmeyer et al., 2009). The continuous absence of the US during CS+ presentations following acquisition leads to extinction learning. While the learning mechanisms of acquisition and extinction have been extensively characterised, far less is known about how threat associations are retrieved and generalised across related stimuli. Theoretically, US expectation is held to be a driver of threat responses during retrieval and generalisation, but the evidence to support this is incomplete.

Fear retrieval, which is the reactivation of stored fear memories when the threat signals (e.g. CS+) are re-encountered, depends on the interaction of memory and salience systems. Additionally, during re-encounters of the CS+, the established memory is not only retrieved, but also updated with new information (Haque et al., 2020). Retrieval-tests are commonly employed as CS+ presentation without a US, which updates the CS-US association with an inhibitory CS-noUS association, in parallel to retrieval (Lonsdorf et al., 2017). Monitoring trial-wise US expectancy might help to disambiguate retrieval from updating processes. The hippocampus has been proposed to be involved during reactivation of associative memory representations based on contextual and perceptual similarity, whereas the insula monitors interoceptive and affective significance, and the PAG coordinates defensive output (Craig, 2009; Milad & Quirk, 2012; Roy et al., 2014). Converging evidence suggests that these regions operate as a dynamic network that infers threat predictions during retrieval of learned associations and updates them when expectations are violated (Menon & Uddin, 2010; Wen et al., 2021). Yet, how these interactions unfold across modalities, i.e. from subjective expectation of the US to physiological arousal and neural responses, is less well understood.

Beyond retrieval of the exact threat predicting stimuli, fear generalisation represents the central process through which learned fear extends to novel but similar stimuli. Behaviourally, this is typically expressed as a graded decrease in US expectancy and arousal as the similarity to the CS+ declines (Dunsmoor & Paz, 2015; Onat & Büchel, 2015; Shepard, 1987). Neuroimaging studies have implicated the hippocampus in representing stimulus similarity, the insula in encoding affective salience and the PAG in scaling the defensive responses (Dunsmoor & Paz, 2015; Mobbs et al., 2009; Onat & Büchel, 2015). However, it remains unclear how these neural computations correspond to subjective gradients of US expectation and physiological indices of generalisation, such as skin conductance or pupil dilation. More specifically, a fundamental question concerns the extent to which neural, behavioural and physiological generalisation responses show a common similarity-based activation pattern.

The present study aims to provide an integrated approach of fear learning, retrieval, and generalisation using a multimodal approach combining behavioural ratings (fear, arousal, US expectancy), autonomic measures (SCR, pupil), and fMRI across two consecutive days. On Day 1, participants acquired threat associations (Acquisition, ACQ). On Day 2, retrieval (RET) and generalisation (GEN) were tested using the original CS+/CS-contingency and a range of generalisation stimuli (GS). We hypothesized that (i) participants would show successful acquisition and retrieval, reflected in increased subjective and physiological responding to CS+ versus CS-, (ii) generalisation would follow a graded, Gaussian-like profile across stimuli, and (iii) neural correlates of these effects would emerge within the amygdala, hippocampus, insula, and PAG, reflecting the integration of memory-based US predictions, interoceptive awareness, and defensive control.

## METHODS

### Participants

We recruited 25 healthy participants between 18 and 40 years (mean age: 23,84) of which 14 were female (∼54%). The participants confirmed to consume less than 15 units of alcohol per week and had not taken illicit drugs in the past two months. Additionally, they showed no contraindications for MRI measurements and had no self-report of any neuropsychiatric diagnoses. All participants gave written, informed consent to participate and received 50€ reimbursement. The study was approved by the local ethics committee and is in line with the Declaration of Helsinki.

### Stimulus Material

For this study, we use a two-day classical fear conditioning paradigm. During testing, visual cues are shown to the participants on a screen using the Presentation® software (Version20.3, Neurobehavioral Systems, Inc., Berkeley, CA, www.neurobs.com) that are either predictive or not predictive of an aversive electrical stimulus (unconditioned stimulus (US)) presented on the participants’ right hand. The visual cues consisted of Gabor patches, i.e. circular sinusoidal gratings convolved by a Gaussian kernel function. The orientation of the conditioned stimuli was 55° and 145°. The assignment of the cues as the US predictive CS+ (and not predictive CS-) was counterbalanced. The electrotactile stimulation to the right hand that served as a US consisted of a train of 3 pulses, each with a duration of 2 ms and an interval of 50ms. The US was delivered via a surface electrode (Specialty Developments, Bexley, UK) on the dorsal surface of the right hand using a DS7A electrical stimulator (Digitimer, Welwyn Garden City, UK). Prior to the habituation, the US intensity was individually adjusted so that it was “painful but tolerable” (mean = 6.52 mA, sd = 3.79 mA, min = 1 mA, max = 15 mA).

The orientation of the Gabor patches can be rotated 180° around the orientation of the CS+, which allowed to construct six generalisation cues (GS) that are orientated according to a circular generalisation gradient around the CS+ in steps of 22.5° (if the CS+ = 55° and the CS-= 145°: GS1 =-12.5°, GS2 = 10°, GS3 = 32.5°, GS4 = 77.5°, GS5 = 100°, GS6 = 122.5°).

In doing so, the responses to the CS+ should always represent the highest peak of the curve, while responses to the conditioned stimulus, which is never followed by a US (CS-) should represent the lowest peak.

### Experimental Procedure

On the first study day, participants underwent the habituation and acquisition (ACQ) trials. During habituation the CS+ and the CS-were shown twice. In this phase, the presentation of the CS+ and CS-was not followed by an US. Then ACQ started, consisting of four blocks of five CS+ and five CS-presentations each (lasting for 6 sec). The CS+ presentations were followed by a US in 80% of the runs (5,5 sec after CS+ onset). Inter-trial intervals (ITI) were shown for 8-10 seconds to account for context-dependent effects. On the second day (24h later), retention (RET) and generalisation (GEN) trials were performed. During RET, five CS+ and five CS-were again presented in four blocks each. This time, none of the CS+ presentations was followed by a US. Afterwards, GEN trials were conducted in which, in addition to the two conditioned stimuli, another six GEN stimuli (GS) were shown whose orientation shifted by 180° around that of the CS+ (see stimulus material, and *Figure 1C*). GEN consisted of two blocks, in which the CS+ and the CS-as well as each of the six GS were presented twice in each block. During the GEN phase, only the first of the CS+ presented was followed by a US to induce a rapid reacquisition of conditioned responses. Otherwise, no stimulus was followed by a US. During ACQ, RET and GEN, the participants rated their expectancy of receiving an US on each trial. Additionally, the participants were asked before and after each of the three test phases of the study how much fear, stress and arousal they felt towards the cues.

**Figure 1.**
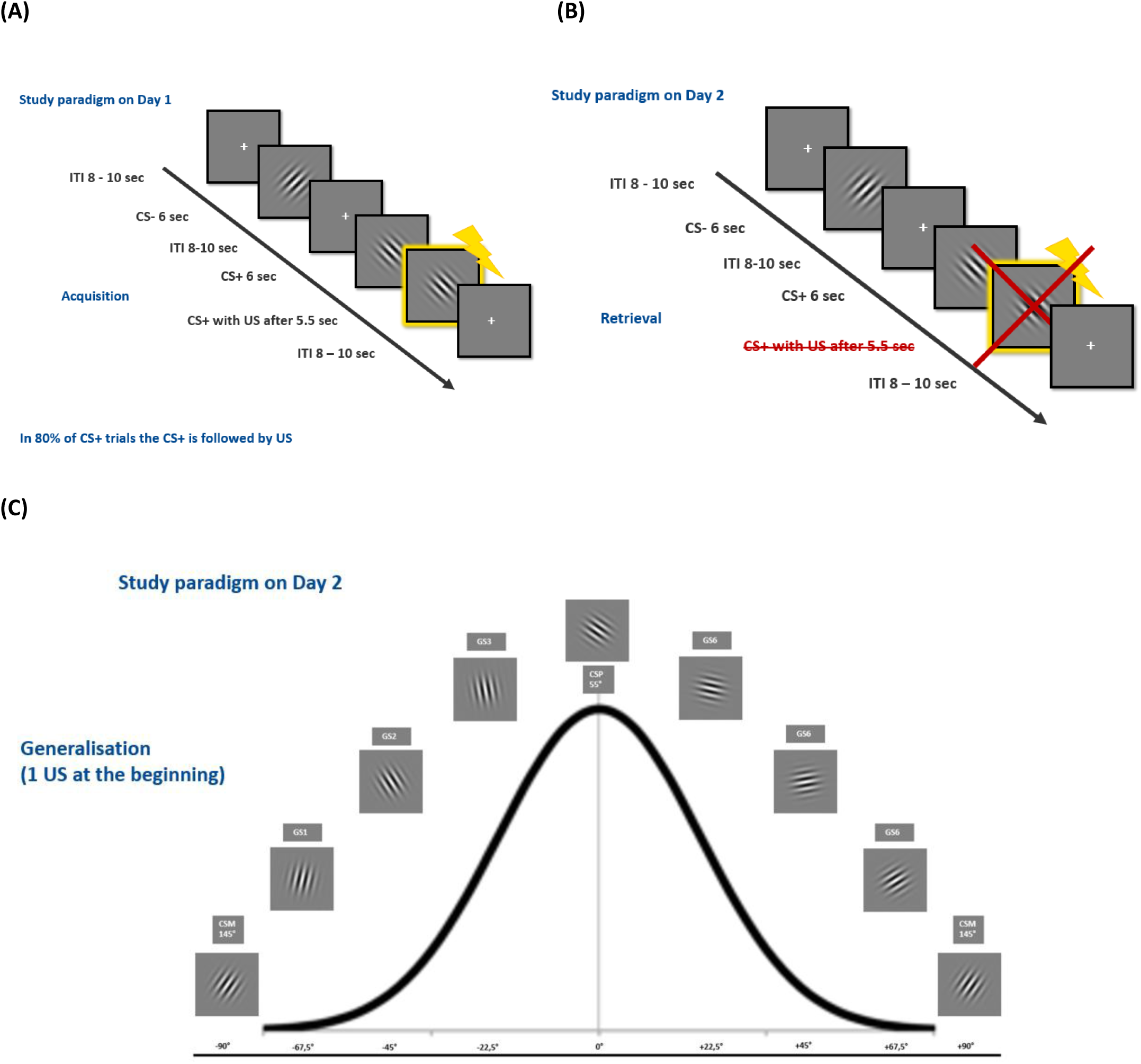
Display of the Fear Conditioning Protocol. **(A)** Acquisition - During the functional MRI scan, participants were shown either a CS+ or a CS-for 6 seconds. In 80% of the CS+ trials the CS+ was followed by a US after 5.5 seconds after CS+ onset. The CS-was never followed by a US. ITIs were shown for 8-10 seconds. **(B)** Retrieval – During the fMRI scan 24 hours later, participants were shown the same CS as the day before, but none of the CS presentations were followed by a US. **(C)** Generalisation – Next to the CSs, participants were shown 6 additional GSs which’s orientation was either close, middle or far away to the orientation of the CS+. Stimuli spanned a continuous orientation gradient from CS-(−90°) through CS+ (0°) back to CS-(+90°). The endpoints were perceptually identical but treated as distinct extremes of a bounded continuum (i.e., not circular).

### Outcome Measures

#### Subjective ratings - Fear, arousal and US expectancy

During ACQ, RET and GEN, the participants rated their expectancy of receiving a US. During the ACQ and RET participants were asked to give a trial-wise binary US expectancy rating (without any visual scale by pressing the upper or lower button on a button box in their right hand (upper button meaning

“Yes, I expect the visual cue to be followed by an electrotactile stimulation” and lower button indicating, “No, I do not expect the visual cue to be followed by an electrotactile stimulation”). During GEN the trial-wise US expectancy was rated on a 10-step Visual Analogue Scale [VAS, 0 (none) –100 (maximally)], which was presented at the onset of each stimulus. Participants had three seconds to rate before the VAS diminished. This was done to verify that the ratings given were rather intuitive. Before and after each of the three test phases of the study the participants were asked, firstly, how much fear and stress and secondly, how much arousal they felt towards each of the stimuli on a VAS from 0 (none) to 100 (very much).

#### SCR data recording

Skin conductance responses were acquired from the hypothenar region of the participant’s left hand with a BIOPAC MP-100 amplifier (BIOPAC® Systems Inc, Goleta, California, USA). For visualisation purposes, the raw data was imported into the Pspm toolbox (version 7.0.0, bachlab.github.io/PsPM) in Matlab (R2022a, The Math Works, Natick, MA, USA) and scored manually. Thereby, data with signal less than 60% or too noisy data was deleted from further analysis.

#### Pupil data acquisition

Pupil diameter data was recorded with an Eyelink 1000 system (EyeLink 1000; SR-Research Ltd., Mississauga, Ontario, Canada) suitable for MRI measurements at a sampling rate of 1000Hz. At the start of each session the eye-tracker was calibrated to record pupil size changes of the right eye.

#### Region of interest

Brain regions of interest (ROI), in which activity pattern changes in memory retrieval are expected to occur were the bilateral amygdala, hippocampus, insula, locus coeruleus (LC) and periaqueductal gray (PAG). These regions were defined as (probalistic) masks of the Harvard-Oxford Atlas for Amygdala, Hippocampus and Insula (Desikan et al., 2006). For the localisation of the LC, a high-resolution, neuro-melanin-sensitive turbo-spin-echo sequence was performed, and a reference mask was then created based on previous findings (Dahl et al., 2019; Dahl et al., 2022). Initially, the PAG ROI was defined by creating a spherical mask (radius: 4mm) centred at the peak voxel coordinates (x = 6, y = 34, z = −6) derived from previous literature (Fairhurst et al., 2007; see also Haaker et al., 2017). However, consistent with our preregistered procedure, analysis of our pilot data identified a distinct peak voxel, prompting the use of a more appropriate PAG mask published by Lojowska et al. (2018).

#### MRI data acquisition

The fMRI data were acquired using a 3 Tesla MR scanner (Prisma, Siemens) with a 64-channel head coil. Each image volume of the T2*-sensitive echo-planar multiband imaging (EPI) sequence (TR = 1.635s, TE = 30ms, flip angle: 70°; 1,5 x 1,5 mm in-plane resolution, field of view = 225mm) consists of 46 axial slices (1.5mm x 1.5mm x 1.5mm with 0.5mm gap). On the second study day, after the RET and GEN phases, high-resolution T1-weighted anatomical brain image (MP-RAGE sequence, 1mm isotropic voxel size, 240 slices) were obtained for normalisation and to aid better coregistration of the functional EPI images. Field maps of the magnetic distortion by physical noise were acquired as well.

### Data Analysis

#### Behavioural data analysis

The data of the fear and arousal ratings before and after ACQ as well as RET and the trial-wise US expectancy ratings were analysed using linear mixed models in R with the lme4 package (Bates et al., 2015).

The fear and arousal ratings were included as the dependent variable, while stimulus (CS+ vs CS-) and time (before vs. after ACQ or RET) were added as factors into the model. For the subject variable since they were tested twice, we added random intercepts (lmer (fear/arousal ∼(1|subject)+stimulus*time)). Afterwards we tested the estimates for the fixed effects, including the interaction with an ANOVA (type 3).

For the analysis of the trial-wise US expectancy ratings for ACQ and RET, each phase was split into four blocks, each consisting of five CS+ trials and five CS-trials. That way we could, as preregistered, focus our analysis on the first block of the RET. The ratings were analysed by adding the US expectancy ratings as the dependent variable, and stimulus and block as factors into a generalised linear mixed model. The subject variable was, again, included as a random intercept (glmer (US expectancy rating * (1|subject)+stimulus*block)) and the fixed effects and their interaction were tested using an ANOVA (type 3) with the car package (Fox & Weisberg, 2018).

The data of the fear and arousal ratings after GEN and the trial-wise US expectancy ratings were analysed similarly, however since this phase consisted of only one block (4 trials for each CS and GS) stimulus was the sole fixed effect term (lmer (fear/arousal/US expectancy ∼(1|subject)+stimulus)). The fixed effect was tested with an ANOVA (type 2), as no interaction was added to the model.

After setting up the models and analysing the ratings, post-hoc tests were used to follow upon significant results from the ANOVAs by comparing the estimated marginal means with the emmeans package (Lenth, 2023) and all results were corrected to account for multiple comparisons (Bonferroni-Holms method).

Additionally, we fitted the trial-wise US expectancy ratings during GEN to a Gaussian model to determine whether the US expectancy follows a Gaussian distribution. For that, we set up the Gaussian function with four parameters:

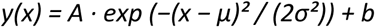

For each participant, the expected Gaussian distribution was set up using the individual peak amplitude (*A* - maximum value of the mean ratings), mean (*μ* - centre of the Gaussian, i.e. stimulus position associated with the peak US expectancy), standard deviation (*σ* - width of the generalisation gradient) and baseline (*b* – accounting for individual differences in overall US expectancy levels) as the initial estimates for the Gaussian parameter. All parameters were averaged across the four repetitions of each stimulus. We then fitted the Gaussian function to the individual US expectancy rating data by using the *lsqcurvefit* function in Matlab, release 2022a (Natick, Massachusetts: The *MathWorks Inc.)*.

#### SCR data analysis

Skin conductance data were analyzed using the PsPM toolbox (version 7.0.0, bachlab.github.io/PsPM), a software package optimized for psychophysiological modelling run in Matlab, release 2022a (Natick, Massachusetts: The *MathWorks Inc.)*. Due to missing data-points (i.e. SCR signal is usable for less than 60%) and noisy data, three data sets for day 1 and five data sets for day 2 were excluded from further analysis. Afterwards, data was trimmed to entail only skin conductance responses during the experimental phases, i.e. ACQ, RET and GEN.

Skin conductance data were analysed by applying a generalised linear model (GLM) as implemented in PsPM. The signal was modelled using the canonical skin conductance response function (SCRF) provided by PsPM with onsets of the experimental conditions as predictors. For analysis of the ACQ data (day 1) each of the conditions (CS+, CS-, US) were included in the design matrix as separate regressors modelling the predicted time courses of experimentally induced activation changes. Parameter estimates for each condition were extracted from the GLM and condition-specific activity was then investigated by computing the main contrast of interest (CS+_ACQ day 1_ > CS-_ACQ day 1_). Similarly, for analysis of the RET data (day 2) each of the conditions (CS+, CS-, US) were included in another design matrix as separate regressors. As we hypothesized that the first block of the RET reflects retention of previous learning, we split the CS presentations into 4 blocks, such that each block contains 5 CS presentations. Then we analysed the mean differential CS responses (CS+ – CS-) by computing the main contrast of interest (CS+_RET1 day 2_ > CS-_RET1 day 2_). Additionally, ACQ data (day 1) were analysed by combining CS+ and CS− trials into a single regressor and modelling trial-wise US expectancy ratings as a parametric modulator. This resulted in two separate regressors, either, a CS of which a following US was expected (CSyes) or not expected (CSno). Both regressors and the US were included in the design matrix to model the predicted time course of the experimentally induced SCR. Parameter estimates for those three conditions were extracted from the GLM and condition-specific SCR were investigated by computing the contrast of interest (CSyes > CSno). For the analysis of the RET data (day 2), we similarly used the trial-wise US expectancy ratings to separate the CS, which resulted in three separate regressors (CSyes, CSno, US) and extracted the parameter estimates from the GLM for further analysis. We hypothesized that on the second day, the SCR is modulated by the US expectancy based on the learned contingencies on day 1. For the analysis of the GEN data (day 2) each of the conditions (CS+, CS-, US and each of the 6 GS) were included in the design matrix as separate regressors.

For the analysis of the SCR data during GEN “stimulus” was the sole fixed effect term entered into a linear mixed effect model (LME) in R (version 4.2.1, www.r-project.org; lmer (fear/arousal/US expectancy ∼(1|subject)+stimulus)). The fixed effect was tested with an ANOVA (type 2), as no interaction was added to the model. Afterwards, post-hoc tests were used to follow upon significant results from the ANOVA by comparing the estimated marginal means with the emmeans package (Lenth, 2023) and all results were corrected to account for multiple comparisons (Bonferroni-Holms method).

Additionally, we applied the same Gaussian modelling approach to the SCR data during the GEN. For each participant, one mean SCR parameter estimate per stimulus (averaged across the four repetitions of each stimulus) was entered into a four parameter Gaussian function:

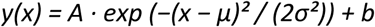

This was then fitted to each participant’s individual GEN gradient using the *lsqcurvefit* function in Matlab, release 2022a (Natick, Massachusetts: The *MathWorks Inc.).* In this model, *A* defines the peak SCR amplitude, *μ* the mean or centre of the Gaussian distribution, *σ* the standard deviation reflecting the width of the GEN gradient and *b* the constant baseline parameter to account for individual differences in overall SCR levels.

#### Pupil size preprocessing and analysis

Pupil dilation data were preprocessed and analysed with the PsPM toolbox (version 7.0.0, bachlab.github.io/PsPM) as implemented in Matlab, release 2022a (Natick, Massachusetts: The *MathWorks Inc.).* Due to technical issues, we only collected pupil dilation data sets of 17 participants. After importing the raw Eyelink data into the toolbox, we followed the default preprocessing pipeline in PsPM. Thereby, the continuous pupil dilation signal was filtered, interpolated and smoothed. To minimise data loss, we increased the maximum gap size for interpolation from 250ms to 1000ms. Missing samples (e.g. due to blinks) were linearly interpolated within a 50ms window before and after the gap. Moreover, in the first-level analysis, the pupil dilation signal was low-pass filtered at 50 Hz to attenuate high-frequency noise and preserve the slower pupil dynamics associated with task-evoked autonomic responses.

In a next step, we applied a GLM as implemented in PsPM. Similarly to analysis of SCR data, we started the analysis of the ACQ data by entering each of the conditions (CS+, CS-, US) into the design matrix as separate regressors. After extracting the parameter estimates for each condition, we computed the main contrasts of interest (CS+_ACQ day 1_ > CS-_ACQ day 1_). For the analysis of the RET data, we also split the CS presentations into 4 blocks (5 trials of each CS per block) to focus the following analysis on the first block of retrieval of previous learning. Then we analysed the mean differential CS responses (CS+ – CS-) by computing the main contrast of interest (CS+_RET1 day 2_ > CS-_RET1 day 2_). Further, for analysis of the ACQ data (day 1) we first included the experimental conditions (CS+, CS-) into one condition (CS+ and CS-combined) and then included the trial-wise US expectancy ratings as a parametric modulator. This either resulted in two separate regressors, a CS of which a following US was expected (CSyes) or not expected (CSno). Both regressors and the US were included in the design matrix to model the predicted time course of the experimentally induced pupil dilation. Parameter estimates for those three conditions were extracted from the GLM and condition-specific pupil sizes were investigated by computing the contrast of interest (CSyes > CSno). For the analysis of the RET data (day 2), we similarly used the trial-wise US expectancy ratings to separate the CS, which resulted in three separate regressors (CSyes, CSno, US) and extracted the parameter estimates from the GLM for further analysis. We hypothesized that on the second day, the pupil size changes are modulated by the US expectancy based on the learned contingencies on day 1. For the analysis of the GEN data (day 2) each of the conditions (CS+, CS-, US and each of the 6 GS) were included in the design matrix as separate regressors.

For analysis of the pupil dilation data during GEN “stimulus” was the sole fixed effect term entered into a linear mixed effect model in R (version 4.2.1, www.r-project.org; lmer (fear/arousal/US expectancy ∼(1|subject)+stimulus)). The fixed effect was tested with an ANOVA (type 2), as no interaction was added to the model. Afterwards, post-hoc tests were used to follow upon significant results from the ANOVA by comparing the estimated marginal means with the emmeans package (Lenth, 2023) and all results were corrected to account for multiple comparisons (Bonferroni-Holms method).

Moreover, as with the US expectancy and SCR data during GEN, we again examined whether the pupil dilation data followed a Gaussian distribution. For each participant, the pupil dilation responses were averaged across the four repetitions per stimulus resulting into one pupil dilation parameter estimate. This was fitted with the same four parameter Gaussian function:

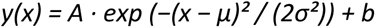

using the *lsqcurvefit* function in Matlab, release 2022a (Natick, Massachusetts: The *MathWorks Inc.).* In this model, *A* defines the peak SCR amplitude, *μ* the mean or centre of the Gaussian distribution, *σ* the standard deviation reflecting the width of the GEN gradient and *b* the constant baseline parameter to account for individual differences in overall pupil dilation levels.

#### fMRI data preprocessing and analysis

The fMRI data were analysed using the SPM12 toolbox (Wellcome Trust Centre for Neuroimaging, London, UK) for MATLAB, release 2021b (Natick, Massachusetts: The *MathWorks Inc.).* In a first step, the first five volumes of each time series were discarded to account for T1 equilibrium effects. Remaining images of the data series were then corrected by the distortion indicated in the field maps.

The fMRI data were then realigned to account for movement of the participants within a mask that was centred on the brainstem. Resulting movement-related distortion were corrected in the following step (unwarping). The further preprocessing included coregistration of the EPI and meanEPI images to the individual high resolution T1 anatomical image. The data were normalised by using the SPM12 DARTEL approach.

Statistical analysis was based on the general linear model as implemented in SPM12. Onsets of the experimental conditions were modelled by a stick function convolved with a hemodynamic response function (HRF). Individual grey and white matter masks, derived from each participant’s anatomical image segmentation, were applied during first-level analyses to restrict estimation to relevant brain regions.

For analysis of the ACQ data (day 1) each of the conditions (CS+, CS-, US, ITI, instructions, button presses) and the intercept were included in the design matrix as separate regressors modelling the predicted time courses of experimentally induced activation changes. The CSs were not grouped into one regressor each, but divided into four blocks. This resulted in four regressors for each, the CS+ and CS-, which covered the four different blocks during ACQ. Additionally, we added 18 physiological regressors to the design matrix to correct for physiological noise. Condition-specific activity was then investigated by computing the main contrast of interest (CS+_day 1_ > CS-_day 1_).

Similarly, for analysis of the RET data (day 2) each of the conditions (CS+, CS-, US, ITI, instructions, button presses), the intercept and 18 physiological noise regressors were included in another design matrix as separate regressors. The CSs were divided into four blocks. As we hypothesized to find a consolidation effect in the first block of the RET, we analysed the mean differential CS responses (CS+ – CS-) by computing the main contrast of interest (CS+_RET1 day 2_ > CS-_RET1 day 2_) for each predefined ROI. Moreover, we tested whether the US expectancy ratings modulated brain activity within our regions of interest. Therefore, we defined all CS (CS+ and CS-) presented during the first block of retention into one regressor and added the mean US expectancy ratings, which were averaged over all subjects, as a parametric modulator (pmod) into the first-level analysis.

Regarding the GEN data, due to technical issues, data from four subjects were unaivailable for the analysis. For the analysis of the GEN data, firstly, we were interested in whether the stimulus gradient correlated with brain activity in voxels of our regions of interest. Therefore, all visual stimuli shown during GEN (CS+, CS- and six GS) were entered into one regressor (REG) next to the remaining conditions (US, ITI, instructions, US expectancy ratings, button presses). Additionally, the trial-wise US expectancy ratings of each subject for each stimulus were added as a parametric modulator (pmod) into the first-level analysis. Secondly, we calculated a full-factorial analysis to investigate whether the stimulus identity (without weighting) might explain variance in activation patterns in different brain regions of interest.

For the second-level group analyses, a mean first-level mask created by averaging individual masks from all subjects was applied, ensuring analyses included only voxels consistently present across subjects. For the factorial designs, the variance assumption was adjusted to “0” assuming equal variances as we have no between-subject factor and a balanced design.

We then investigated the time-course and interactions of the condition-specific activity. For that we set up a full factorial design with the two factors stimulus and block (2×4 design) and computing the main contrasts of interest over all blocks (ACQ; CS+ > CS-) and per block (RET; CS+ > CS-block 1, CS+ > CS-block 2, CS+ > CS-block 3, CS+ > CS-block 4).

For the parametric modulation analysis during the RET phase, a second-level group analysis was performed using the contrast images generated from each subject’s first-level analysis. Specifically, individual contrast images reflecting the parametric modulation of CS – related activity by mean US expectancy ratings were entered into a random-effects one-sample t-test at the second level.

Moreover, we assessed functional connectivity using psychophysiological (PPI) analyses in SPM12. Specifically, we restricted the connectivity analysis to the first block of RET. The seed ROI was defined as a 6-mm-radius spherical mask centred on the group-level peak voxel within the left hippocampus identified in the second-level-analysis of the contrast CS+ > CS-block 1 (MNI: −30 −12 - 22). The mask was created in SPM12 and used to extract the seed time series for PPI. For each participant, the PPI term was entered into a first-level GLM that also included motion parameters. The resulting contrast estimates were then submitted to a second-level group analysis to test for differences in stimulus discrimination.

For the GEN data, a separate full factorial analysis was conducted to investigate whether stimulus identity explained significant variance in activation patterns across predefined regions of interest. The variance was also set to “0” assuming, again, equal variances. Similarly to RET, first-level contrast images from the GEN phase, representing trial-wise modulation of neural activity by subject-specific US expectancy ratings, were entered into another second-level random-effects analysis. These second-level analyses allowed for assessment of consistency and generalizability of the parametric modulation effects and stimulus-related activations across the group.

The resulting estimates were spatially smoothed (4mm) to account for individual differences in spatial location of the activation before adding them to the second level.

Family-wise-error correction (FWE) was used to account for multiple comparisons. The significance level was set at *P* <.05 and *P* – values are reported at peak level.

## RESULTS

**Table 1.** Overview of the sample characteristics.

| Sample characteristics |  |
| --- | --- |
| **Age (Mean / SD)** | 23.84 / 3.44 |
| **Female / Male** | 14 / 11 |

### Behavioural results

In accordance with successful threat learning, our analysis of the fear, arousal and trial-wise US expectancy ratings showed that participants differentiate more strongly between CS+ and CS-after ACQ than before ACQ. Specifically, ANOVAs for fear, arousal ratings and US expectancy ratings revealed a significant stimulus-by-time interaction for day 1 (fear ratings: *F*(1, 72) = 10.75, *p* = .001; arousal ratings: *F*(1, 72) = 6.66, *p* = .011; US expectancy ratings: *F*(1, 953.6) = 9.18, *p* < .001). Larger CS differentiation was found after ACQ as compared to before (fear ratings: *t*(52.45) = 2.26, *p* = .028; arousal ratings: *t*(67.59) = 2.04, *p* = .045; US expectancy ratings: *t*(218) = 4.31, *p* < .001). For day 2, the analysis of the US expectancy revealed that subjects retrieved the learned association from day 1 in the first block of RET. As such, analyses indicated larger US expectancy ratings to the CS+ as compared to the CS-(stimulus main effect: *F*(1, 968) = 42.74, *p* < .0001). This discrimination between CS+ and CS-in the first block decreased towards the last block of RET indicated by a stimulus*block interaction (*F*(3, 968) = 10.21, *p* < .0001; post-hoc *t*(57.26) = −2.94, *p* = .005). Similarly, the analysis of fear and arousal ratings revealed a stimulus-by-time interaction (fear ratings: *F*(1, 72) = 7.77, *p* = .007; arousal ratings: *F*(1, 72) = 4.81, *p* = .031). That is, the discrimination between CS+ and CS-was larger at the beginning of RET, as compared to the end of RET for the fear ratings (*t*(59.08) = −2.06, *p* = .044), whereas this comparison did no yield statistical significance for the arousal ratings (*t*(58.6 6) = −1.61, *p* = .112).Results are summarized in *Figure 2*.

**Figure 2.**
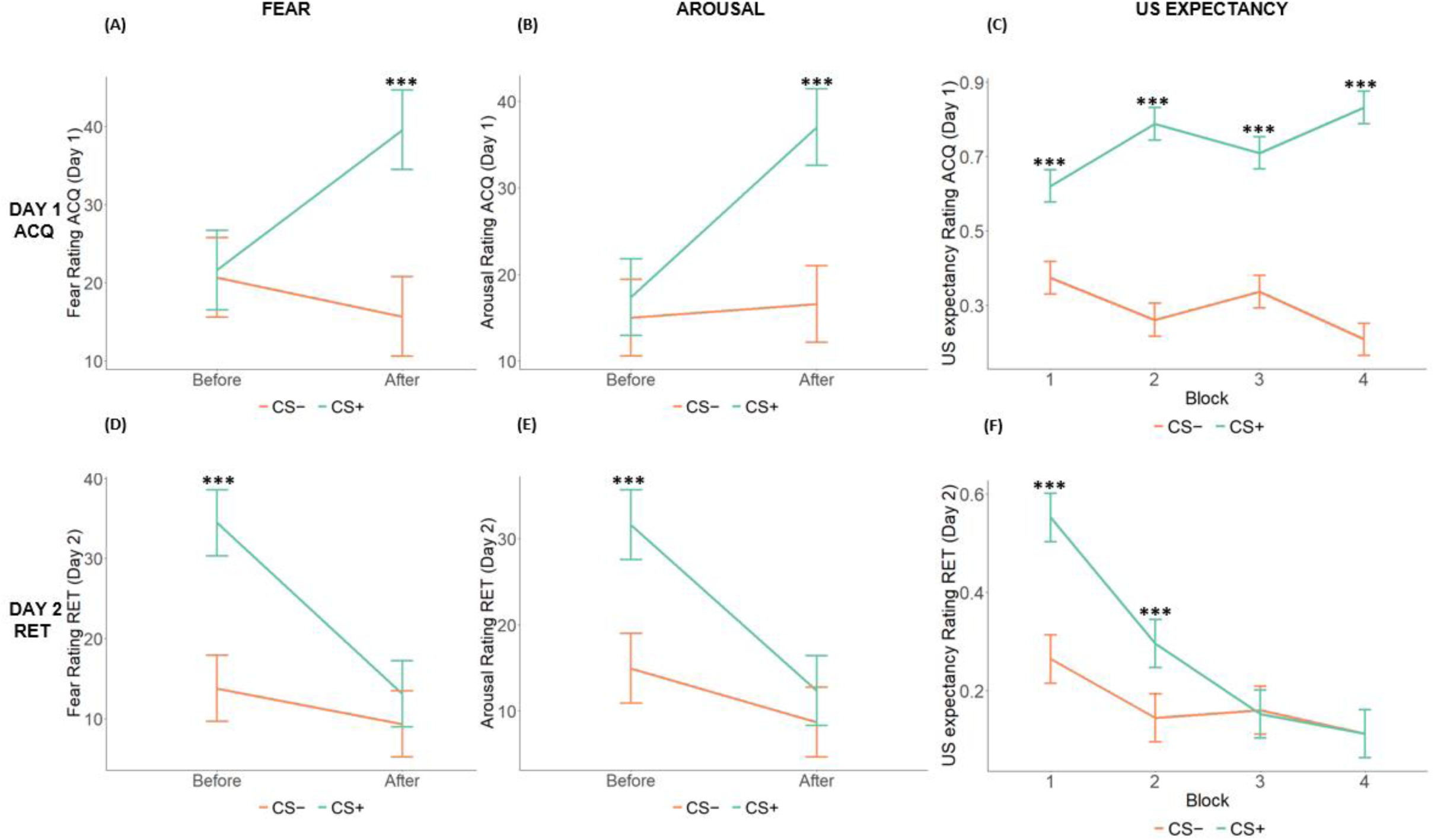
Mean subjective ratings show differential responding towards end of ACQ and in the beginning of RET. Mean subjective ratings of fear **(A, D)**, arousal **(B, E)**, and US expectancy **(C, F)** during ACQ (Day 1) and RET (Day 2). Ratings are shown separately for the conditioned stimuli (CS+ and CS-). Participants showed increased ratings for CS+ than CS-trials after ACQ, indicating successful differential learning. This differentiation between CS+ and CS-was still present on Day 2 when starting RET and decreased towards the end of the RET. *Note:* Fear and arousal ratings were rated before and after ACQ and RET, while US expectancy were trial-wise ratings during ACQ and RET. Error bars represent ± 1 SEM. *Abbrevations:* ACQ = acquisition, RET = retrieval, CS = conditioned stimuli. *** *P* ≤.001

The analyses of the GEN further revealed generalisation of fear, arousal and US expectancy. The stimulus effect in the ANOVA of US expectancy indicated that the orientation of stimuli changes the US expectancy (ANOVA:stim (*F*(8, 867) = 12.042, *p* < .0001) and feelings of fear and arousal (fear ratings: (*F*(8, 192) = 1.72, *p* = .097; arousal ratings: (*F*(8, 192) = 2.38, *p* = .018). Results are summarized in *Figure 3*.

**Figure 3.**
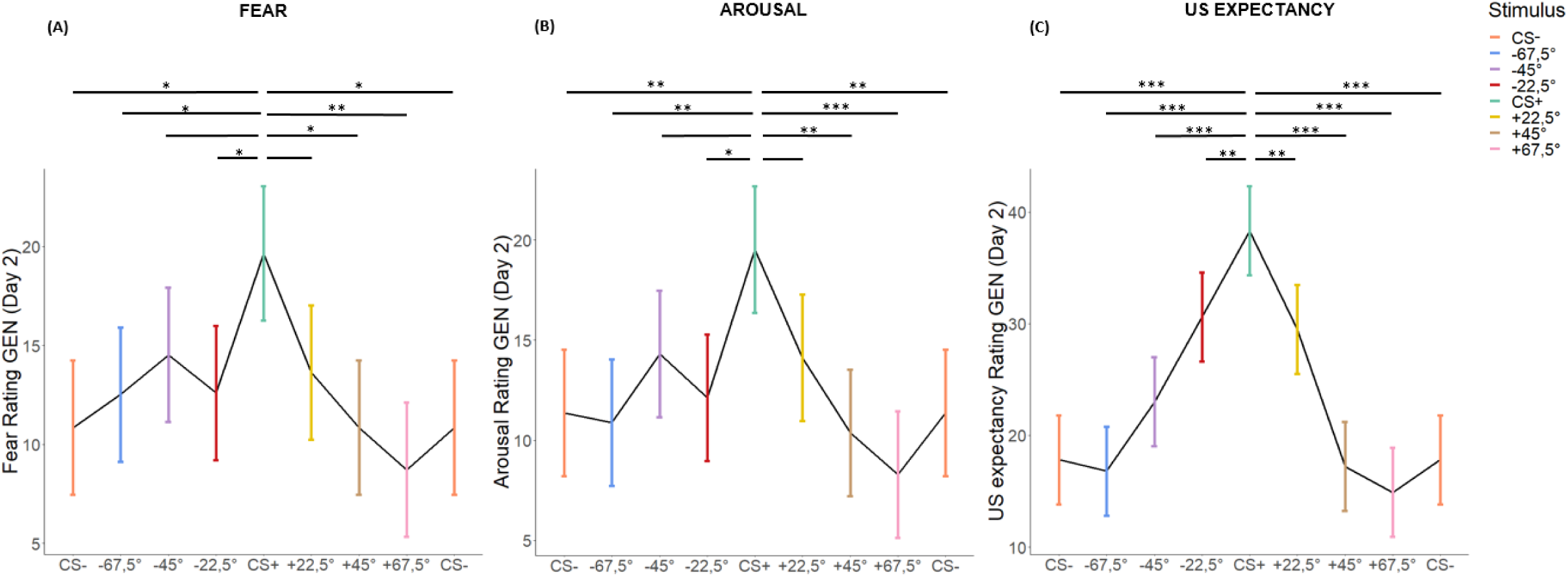
Mean subjective ratings show generalisation pattern. Mean subjective ratings of fear **(A)**, arousal **(B)**, and US expectancy **(C)** during GEN (Day 2). Ratings are shown separately for the conditioned stimuli (CS+ and CS-) and six GEN stimuli. Participants showed peak ratings for CS+, which decreased towards the most dissimilar stimuli. *Note:* Fear and arousal ratings were rated after GEN, while US expectancy were trial-wise ratings during GEN. Error bars represent ± 1 SEM. *Abbrevations:* GEN = generalisation, CS = conditioned stimuli. \**P* ≤.05 \*\**P* ≤.01 *** *P* ≤.001

To examine whether US expectancy generalised according to a Gaussian distribution across the stimuli, we fitted a four parameter Gaussian function to individual US expectancy ratings during GEN. The model captured the individual data well (median R^2^ = .84), although cross-validated predictive performance was modest (median Q^2^ = - 0.45). Using permutation-based significance testing on Q2, 6 out of 25 participants (24%) showed a significant Gaussian fit after controlling for multiple comparisons (FDR < .05). Across participants, the peak amplitude of the GEN gradient was A = 37.03 ± 25.90. The centre of the curve was located at a mean of *μ* = −20.23 ± 41.76°, indicating that the highest US expectancy was on average shifted towards stimuli slightly away from the CS+. The width of the gradient was *σ* = 19.48° ± 11.65°, reflecting a moderately narrow GEN profile (see *Figure 4*). Taken together, these results suggest that US expectancy during GEN shows a partially Gaussian-like pattern at the group level (see *Figure 4*).

**Figure 4.**
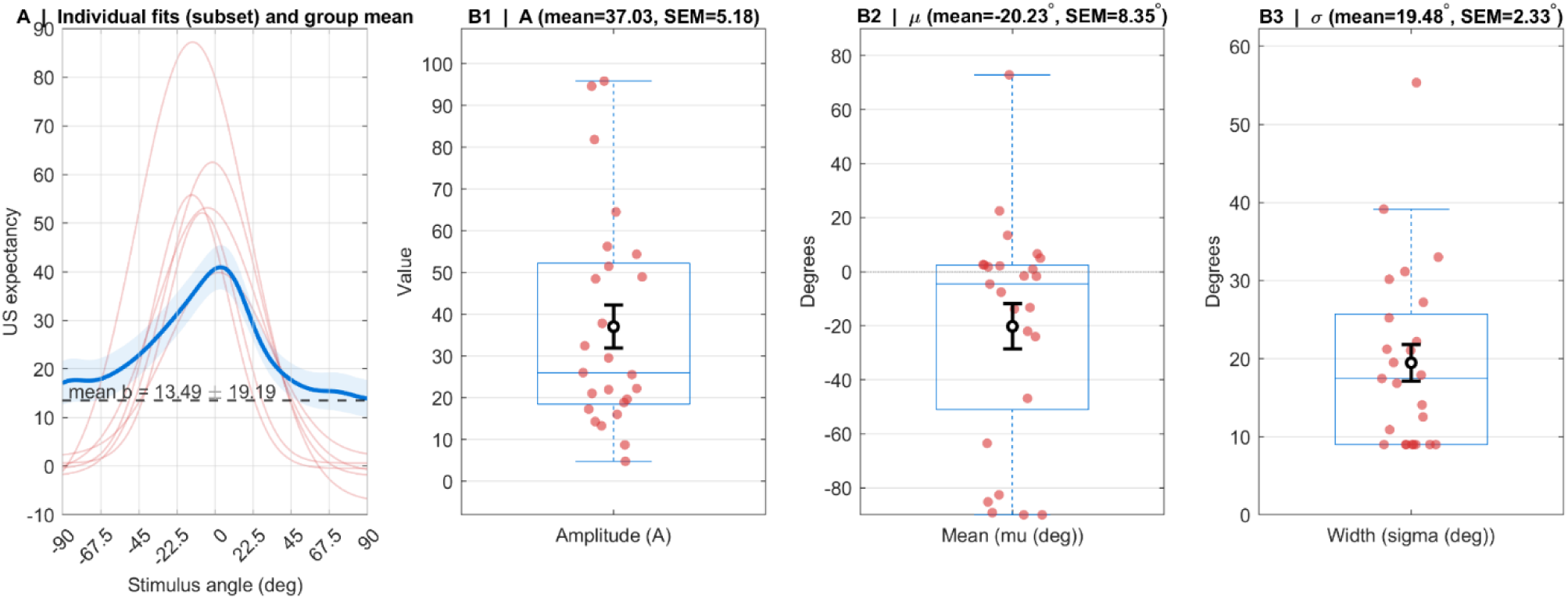
Gaussian model fits of US expectancy ratings. **(A)** Individual participant fits (thin red lines) and the group mean fit (thick blue line) across the six GEN stimuli (orientations ranging from −90° to +90°). The Gaussian model captured a peak around the CS+ (0°), indicating a graded generalisation of US expectancy. **(B1 – B3)** Distributions of individual parameter estimates: amplitude **(B1)**, mean **(B2)**, and width **(B3)**. Black circles represent the group mean ± SEM. On average, the fitted curves show a positive amplitude, a peak slightly shifted towards the left, and a moderate width, indicating a moderately narrow generalisation around the CS+.

### SCR results

For the analysis of SCR data of the ACQ (day 1) we had to exclude data of three subjects (N = 22) on day 1 (ACQ) and six subjects (N = 19) on day 2 (RET), because the recorded data was too noisy. In the subsequent group-analysis, we firstly tested whether SCR increases more towards CS+ presentations than CS- presentations. A one-sample t-test revealed no difference in responses to either CS (*t*(21) = .783, *p* = .443) during ACQ. Additionally, testing the differential responses during block 1 of RET, we did not find any difference between CS+ and CS-(*t*(18) = .901, *p* = .381).

The second analysis was motivated by the assumption that the SCR might be more sensitive to trial-wise US expectancy ratings modulating the activation pattern than a classical CS+ > CS-contrast, as this approach captures expectancy-based arousal during learning and retrieval. On day 1 (ACQ) the analysis revealed no difference in responses to CS-trials in which participants expected an US (CSyes) compared to CS-trials in which participants expected no US (CSno; CSyes > CSno; (*t*(21) = 1.6, *p* = .123; see *Figure 6A*). This shows that the expectation of the US did not significantly modulate the SCR during ACQ. On day 2 (RET) the group-analysis based on US-expectation revealed higher SCR during CS-trials in which an US was expected (CSyes) compared to trials in which no US was expected (CSno; CSyes > CSno; *t*(18) = 3.55, *p* = .002; see *Figure 6A*), showing that SCR is tightly linked to US expectations during retrieval of threat associations.

**Figure 5.**
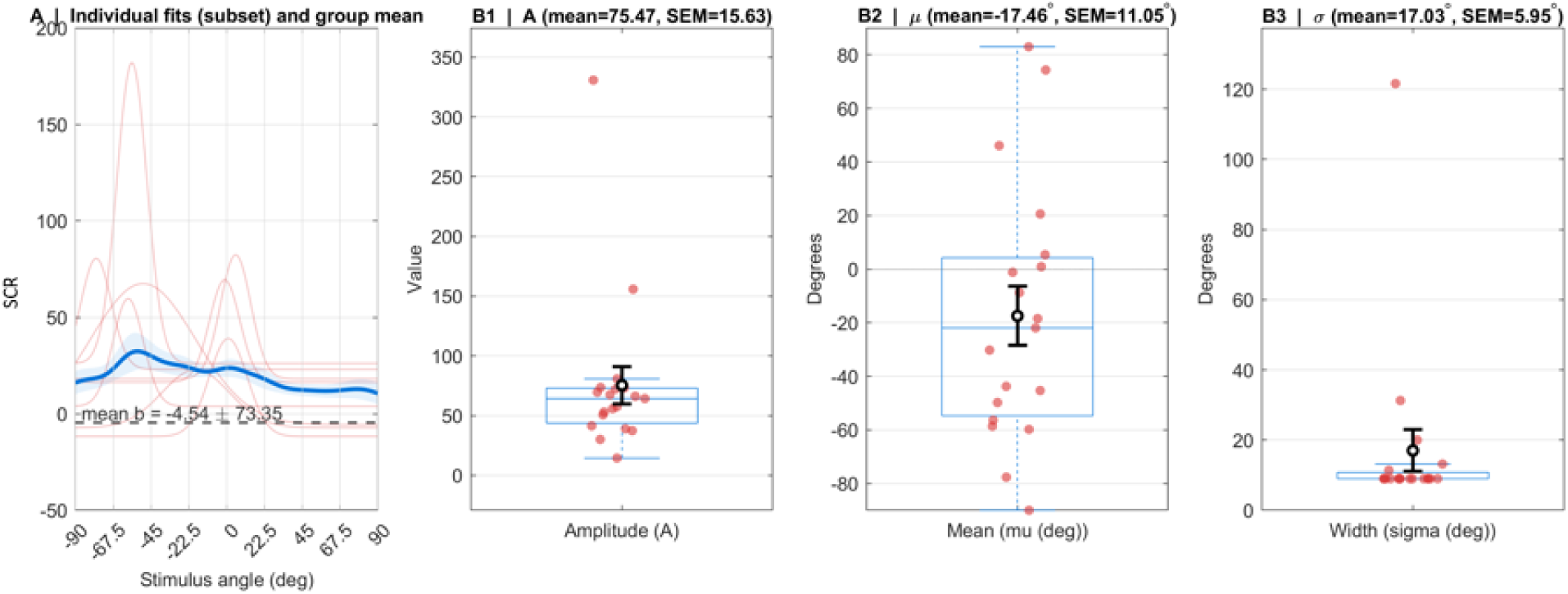
Gaussian model fits of SCR. **(A)** Individual participant fits (thin red lines) and the group mean fit (thick blue line) across the six GEN stimuli (orientations ranging from −90° to +90°). The Gaussian model provided a modest fit to the SCR data, with no clear peak around the CS+ (0°) and high variability. **(B1 – B3)** Distributions of individual parameter estimates: amplitude **(B1)**, mean **(B2)**, and width **(B3)**. Black circles represent the group mean ± SEM. SCR estimates should therefore be interpreted with caution, as the model did not capture the underlying generalisation pattern.

**Figure 6.**
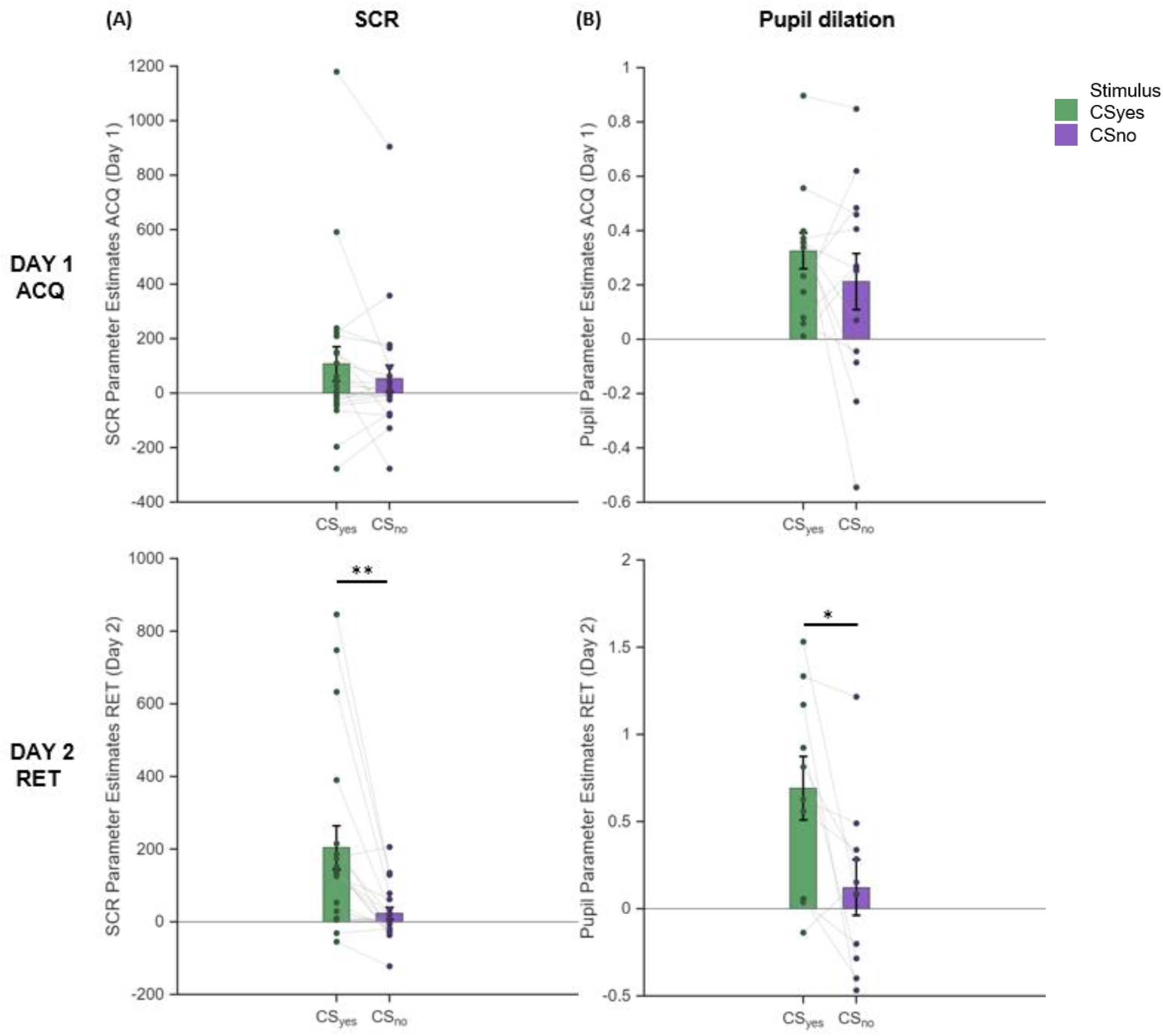
US expectancy driven arousal pronounced in RET but not ACQ in SCR and Pupil dilation. Mean SCR **(A)** and pupil dilation **(B)** parameter estimates during ACQ (Day 1, first row) and RET (Day 2, second row) for CS trials in which the US was expected (CSyes) or not expected (CSno). During ACQ, no differential physiological responses were found between the CS. During RET, both SCR and pupil dilation estimates were significantly higher for CSyes than CSno, indicating physiological diffentiation between the CSs based on US expectancy. *Abbrevations*: SCR = Skin Conductance Response, ACQ = acquisition, RET = retrieval, CS = conditioned stimuli. \**P* ≤.05 \*\**P* ≤.01

The analyses of the GEN further revealed generalisation of the SCR. The stimulus effect in the ANOVA indicated that SCR changed with the orientation of the stimuli (ANOVA: stim (*F*(8, 144) = 3.37, *p* = .001). Regarding the fitting of the individual SCR data to the Gaussian function, we found a modest fit (median R^2^ = .49) and low cross-validated predictive performance (median Q^2^ = −2.39). After controlling for multiple comparisons, none of the participants’ SCR data showed a significant Gaussian fit. The peak amplitude of the GEN gradient was A = 75.47 ± 68.13. The centre of the curve was located at a mean of −17.46° ± 48.18°, indicating that the highest response shifted slightly towards the left. The width of the generalisation gradient was *σ* = 17.03° ± 25.93°, indicating a moderately narrow GEN profile. Overall, this shows that the SCR data does not fit a Gaussian model on the group-level (see *Figure 5*).

### Pupil dilation results

After preprocessing, we had to exclude four subjects from further analysis of ACQ data (day 1, N = 13) and another three subjects from further analysis of RET data (day 2, N = 10), because the data was too noisy. Firstly, we tested whether the pupil dilates more towards CS+ presentations than CS-presentations. A one-sample t-test revealed a significant difference when contrasting CS+ > CS-trials (*t*(12) = 3.669, *p* = .003). When contrasting CS+ against CS-trials during the first block of RET, we did not find any differences in pupil dilation (*t*(9) = .610, *p* = .559).

Secondly, we followed the same US expectancy based approach as in the analysis of the SCR data. To do that, we tested whether pupil dilation during ACQ follows the individual expectancy of receiving a US when being confronted with the CS+ or the CS-. When contrasting pupil dilation size to CS-trials in which participants expected an US with CS-trials in which participants expected no US (CSyes > CSno) we did not find any difference in pupil size (*t*(12) = 1.057, *p* = .311, see *Figure 6B*). For the analysis of RET (day 2), the result of the one-sample t-test showed higher pupil dilation responses to CS-trials in which participants expected an US than to CS-trials (CSyes) in which they not expected an US (CSno; CSyes > CSno; *t*(9) = 2.817, *p* = .020; see *Figure 6B*).

The analyses of the GEN further revealed generalisation of pupil dilation responses. The stimulus effect in the ANOVA indicated that pupil dilation changed with the orientation of the stimuli (ANOVA: stim (*F*(8, 64) = 3.64, *p* = .002). Modelling the pupil dilation data to the Gaussian function showed a good fit (median R^2^ = .735), however, cross-validated predictive performance was modest (median Q^2^ = - 0.73). After controlling for multiple comparisons, none of the participants’ pupil dilation data showed a significant Gaussian fit. Across participants, the peak amplitude of the GEN gradient was A = 0.95 ± 0.64. The centre of the curve was located at a mean of *μ* = −7.91 ± 23.15°, indicating only a slight shift away from the CS+. The width of the gradient was *σ* = 30.48° ± 34.20°, reflecting a wide generalisation profile. These results suggest a good fit of the sample’s pupil dilation data to the Gaussian model, however, with high inter-individual variability (see *Figure 7*).

**Figure 7.**
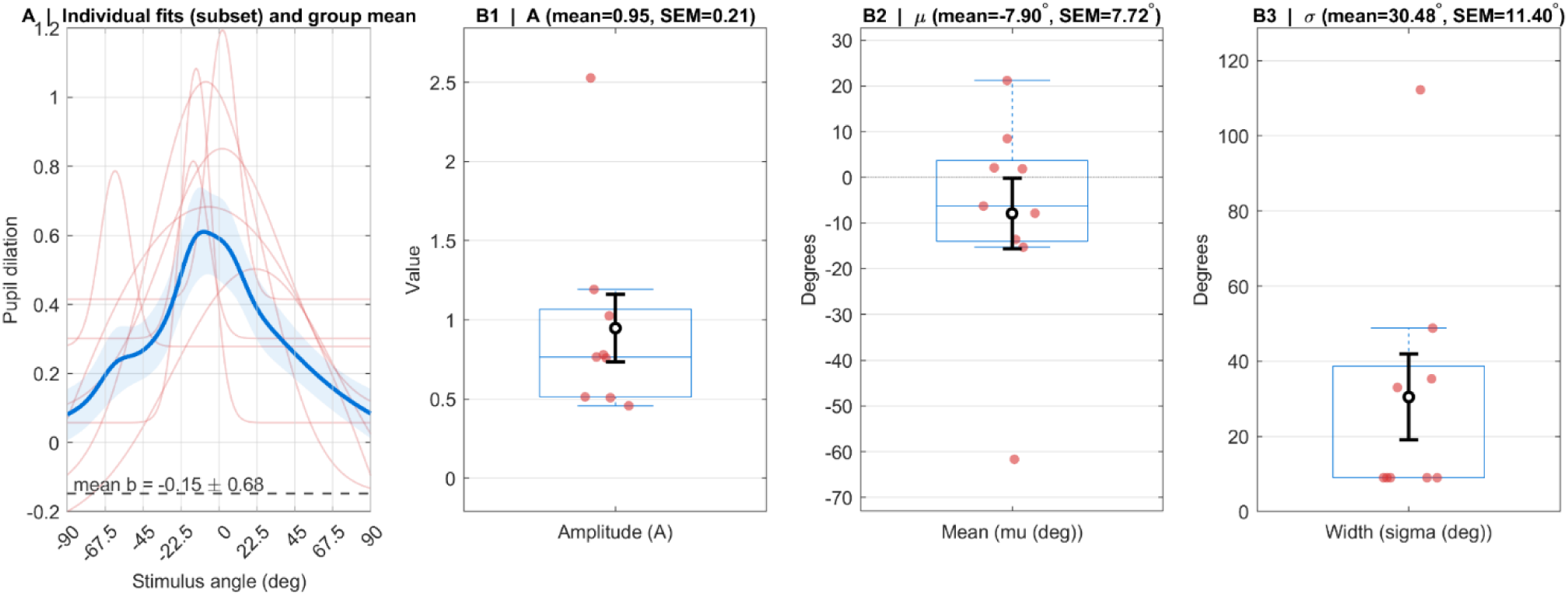
Gaussian model fits of Pupil dilation. **(A)** Individual participant fits (thin red lines) and the group mean fit (thick blue line) across the six GEN stimuli (orientations ranging from −90° to +90°). The Gaussian model showed a good fit to the pupil dilation data, with responses peaking around the CS+ (0°). **(B1 – B3)** Distributions of individual parameter estimates: amplitude **(B1)**, mean **(B2)**, and width **(B3)**. Black circles represent the group mean ± SEM. On average, the fits show a moderate amplitude, a peak slightly shifted towards the left and a broad width, suggesting moderately generalised pupil responses around the CS+.

### fMRI results

First, fMRI data of the ACQ (day 1) was analysed. When contrasting CS+_day 1_ > CS-_day 1_, the full factorial design analysis revealed activation in bilateral insula (Peak Montreal Neurological Institute coordinates (x, y, z): −34 9 6; *t*(24) = 4.27, *p*[pFWE] *=* .007 (left insula), 39 9 2; *t*(24) = 4.67, *p*FWE = .002 (right insula)) and the left PAG ((−2 −30 −4; *t*(24) = 4.10, *p*FWE = .003), see *Table 2* and *Figure 8*).

**Figure 8.**
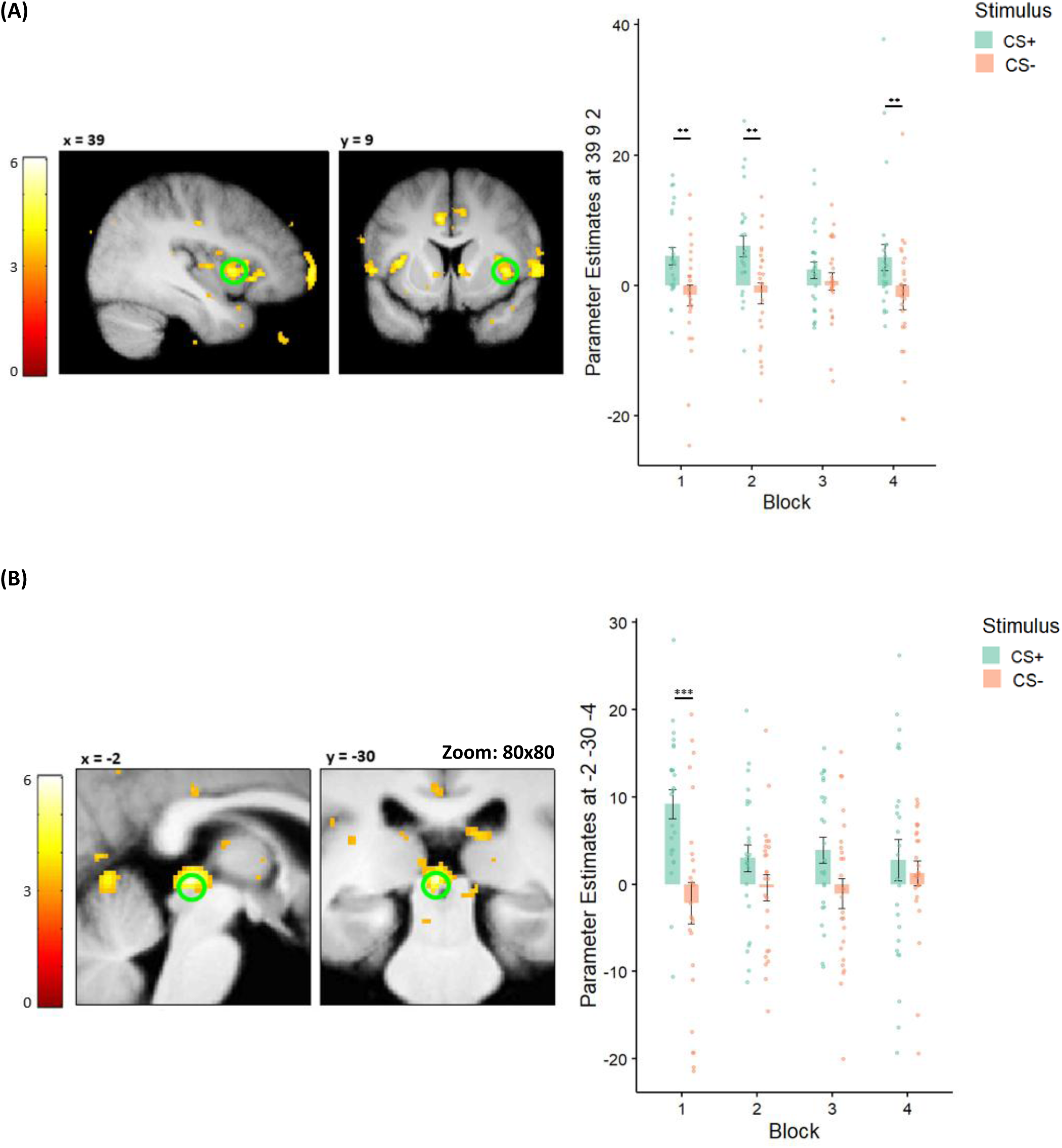
fMRI results in (A) right insula and (B) PAG during ACQ. During ACQ participants show significantly higher BOLD activity for CS+ (green) than CS-(orange) in the **(A)** right insula in block 1, 2 and 4, and **(B)** in the first block of ACQ in the left PAG, indicating successful neural differentiation between CS+ and CS-. *Abbreviations:* CS = conditioned stimulus, PAG = Periaqueductal Gray. \*\**P* ≤.01 *** *P* ≤.001

**Table 2.** Results of full factorial analysis (DAY 1 – ACQ), N = 25.

| Region of interest | Hemisphere | t-score | MNI coordinates |  |  | P FWE-corr |
| --- | --- | --- | --- | --- | --- | --- |
|  |  |  | X | Y | Z |  |
| CS+>CS- |  | df = 24 |  |  |  |  |
| Amygdala | L | - | - | - | - | - |
|  | R | - | - | - | - | - |
| Hippocampus | L | - | - | - | - | - |
|  | R | 3.35 | 27 | -39 | 2 | .130 |
| Insula | L | 4.27 | -34 | 9 | 6 | .007** |
|  | R | 4.67 | 39 | 9 | 2 | .002** |
| PAG | L | 4.10 | -2 | -30 | -4 | .003** |
|  | R | - | - | - | - | - |
| LC | L | - | - | - | - | - |
|  | R | - | - | - | - | - |

*Note:* P-values are reported at peak level (family-wise-error-corrected). *Abbreviations:* L = left, R = right, PAG = Periaqueductal Gray, LC = Locus Coeruleus.
\*\* $P \leq .01$

Secondly, fMRI data of RET (day 2) was analysed to examine brain regions that reflect the retention of the learned associations from day 1. We found higher activation in the bilateral hippocampus for CS+ trials than CS-trials in the first block of RET (left hippocampus: −30 −12 −22; *t*(24) = 4.19, *p*FWE = .009; right hippocampus: 28 −32 −9; *t*(24) = 3.70, *p*FWE = .047). Additionally, we found higher activation in the left insula (−38 −2 10; *t*(24) = 3.71, *p*FWE = .046) and in the right PAG (2 −28 - 10; *t* (24) = 3.59, *p*FWE = .013; see *Table 3* and *Figure 9*). We were furthermore interested in whether the trial-wise US expectancy ratings modulate the activity in our regions of interest, corresponding to the physiological measures of arousal during RET. We found that the trial-wise US expectancy to the CSs was reflected by activity in the left insula (−39 −4 −8; *t*(24) = 5.52, *p*FWE = .005) and right PAG (2 - 28 −10; *t*(24) = 3.91, *p*FWE = .025) as well as trend-wise in the left hippocampus (−33 −30 −12; *t*(24) = 4.18, *p*FWE = .068, see *Table 4* and *Figure 10*). To further characterize the functional interaction of the left hippocampus during RET, we tested the connectivity to other regions of interest. To do that we set up a psycho-physiological interaction. Results showed increased functional connectivity during the first block of retrieval between the left hippocampus and bilateral insula (−38 −6 −4; *t*(24) = 4.75, *p*FWE = .025 (left insula), 39 3 4; *t*(24) = 4.45, *p*FWE = .047 (right insula), see *Table 5* and *Figure 11*).

**Figure 9.**
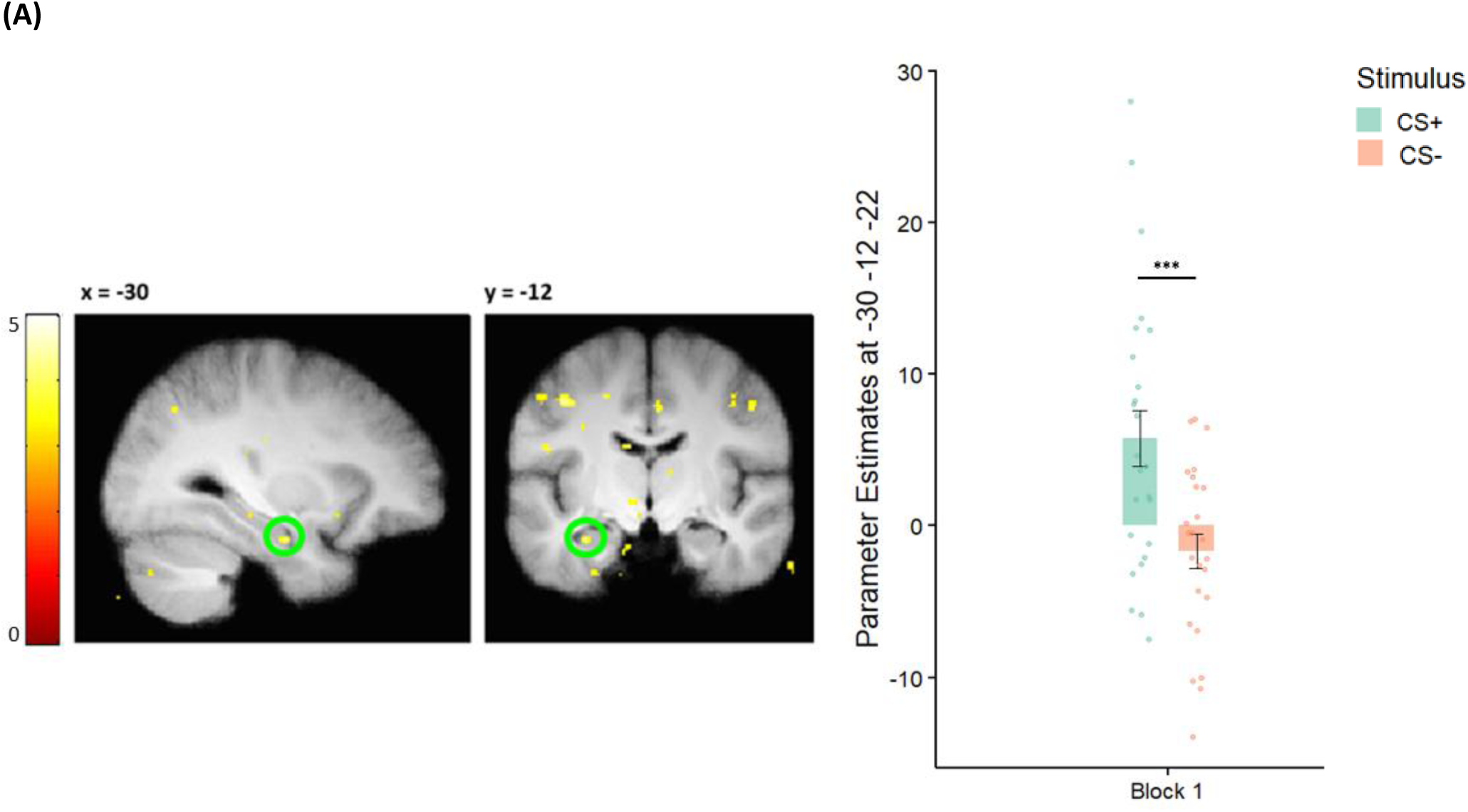
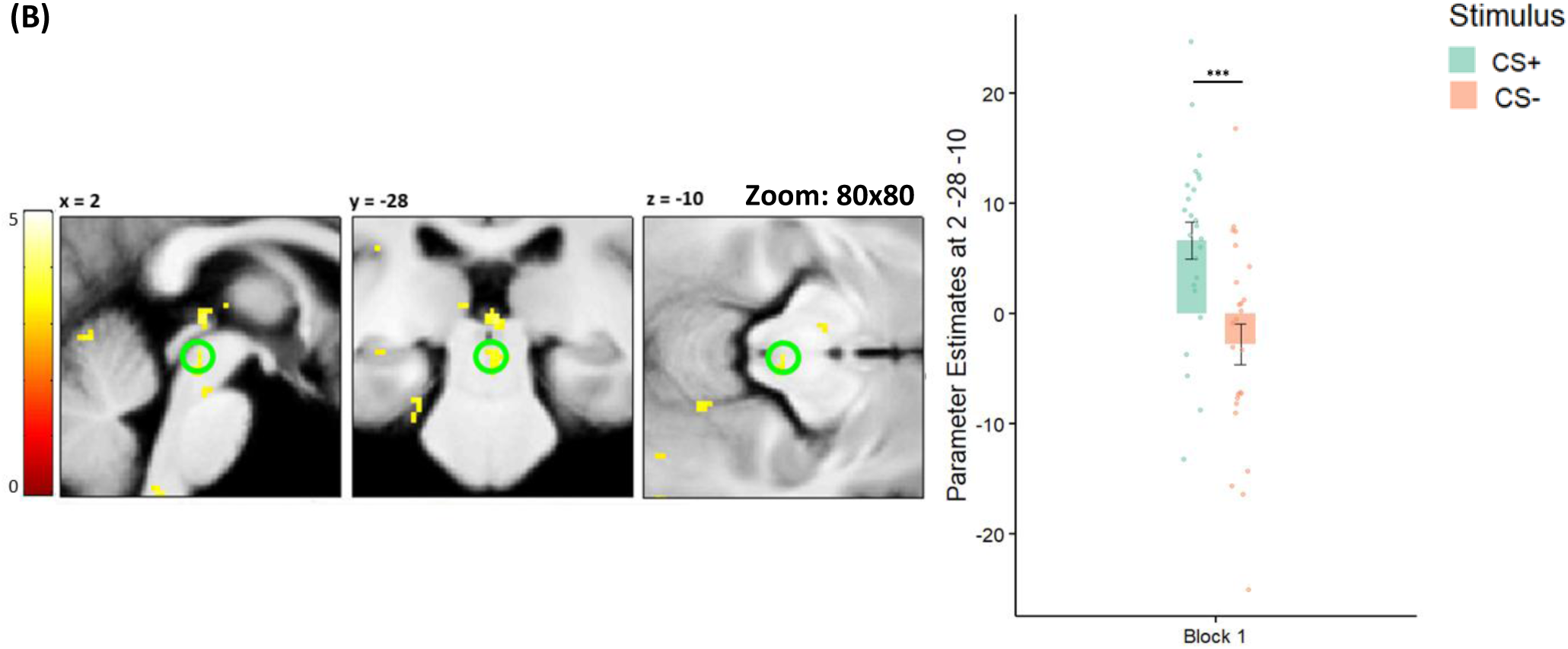
fMRI results in (A) left hippocampus (B) PAG during first block of RET. During the first block of the RET, participants show higher BOLD activity in the **(A)** left hippocampus and **(B)** PAG for CS+ (green) than CS (orange), indicating successful retrieval of the CS contingency. *Abbreviations:* CS = conditioned stimulus, PAG = Periaqueductal Gray. \*\**P* ≤.01 *** *P* ≤.001

**Figure 10.**
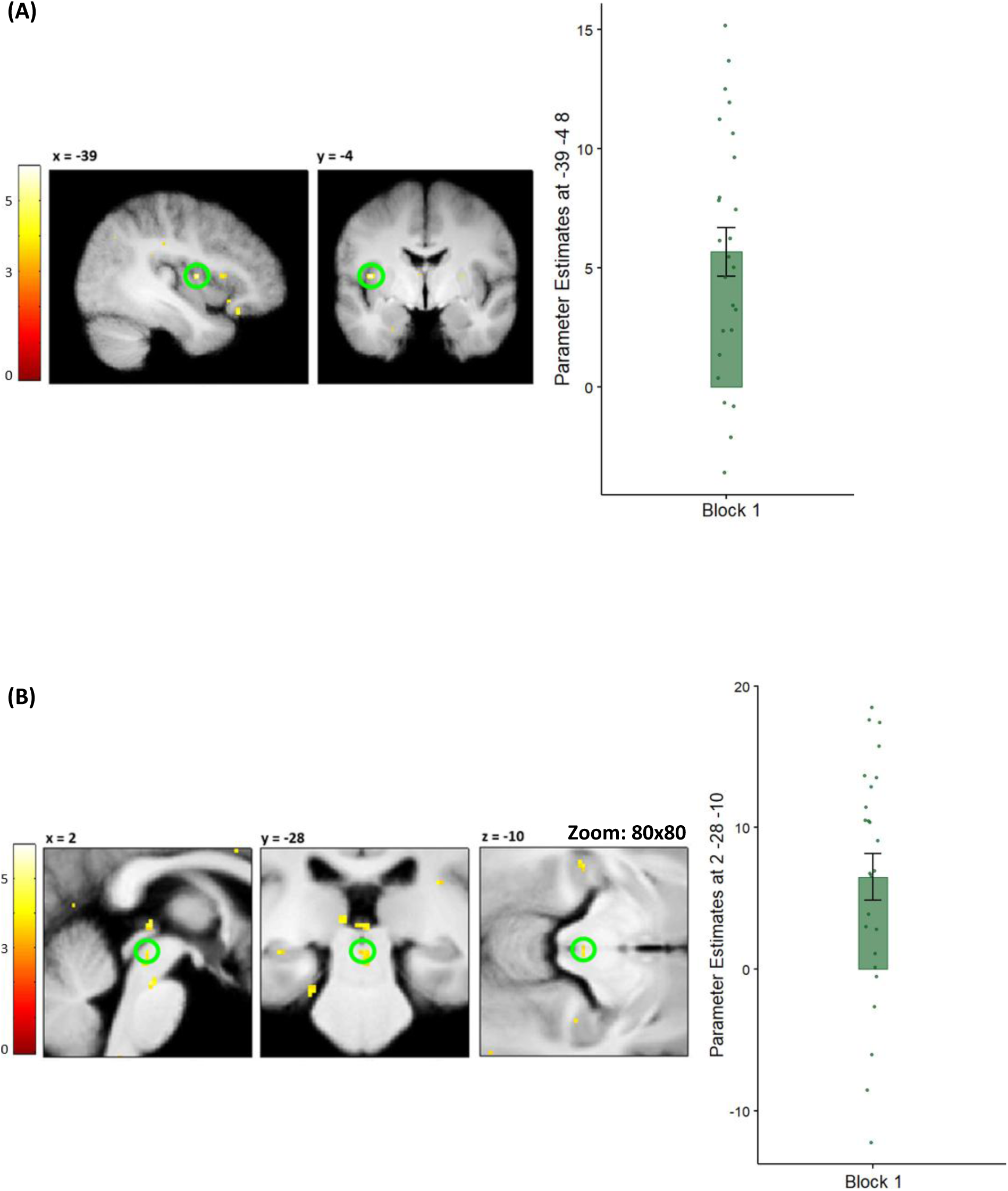
US expectancy driven modulation of BOLD responses during RET in (A) left insula and (B) PAG. Results of the parametric modulation analysis show that during the first block of RET the US expectancy significantly modulates BOLD activity in the **(A)** left insula and **(B)** PAG. *Abbreviations:* RET = retrieval, PAG = Periaqueductal Gray.

**Figure 11.**
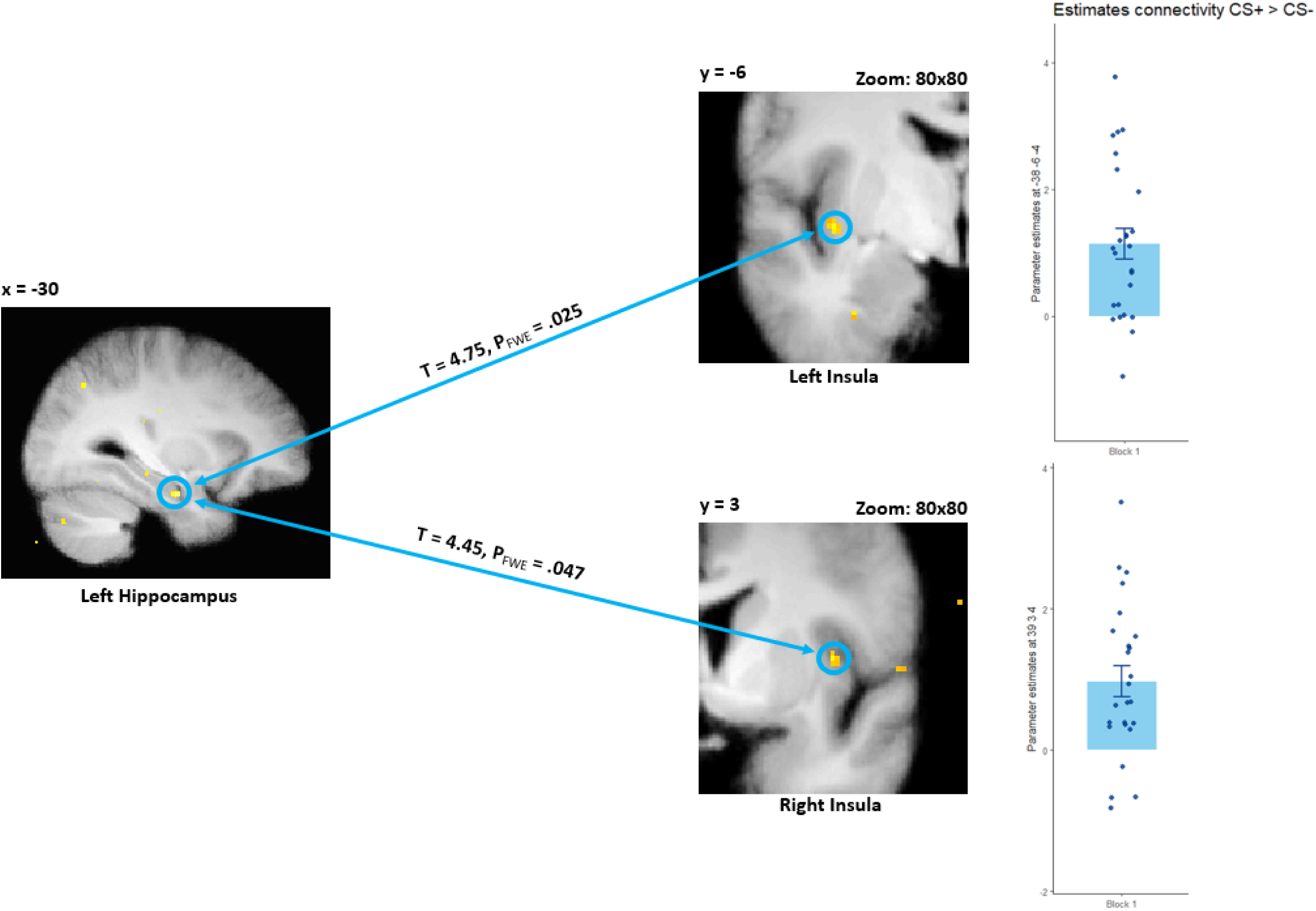
Results of physio-psychological interaction analysis during the first block of RET. Results of the physio-psychological interaction analysis show that during the first block of RET the left hippocampus has significantly increased connectivity to bilateral insula when contrasting CS+ against CS-. *Note: P*-values are reported at peak level (family-wise-error-corrected). *Abbrevations:* RET = retrieval.

**Table 3.** Results of full factorial analysis (DAY 2 – RET 1), N = 25.

| Region of interest | Hemisphere | t-score | MNI coordinates |  |  | P FWE-corr |
| --- | --- | --- | --- | --- | --- | --- |
|  |  |  | X | Y | Z |  |
| CS+ret1>CS-ret1 |  | df = 24 |  |  |  |  |
| Amygdala | L | - | - | - | - | - |
|  | R | - | - | - | - | - |
| Hippocampus | L | 4.19 | -30 | -12 | -22 | .009** |
|  | R | 3.70 | 28 | -32 | -9 | .047* |
| Insula | L | 3.71 | -38 | -2 | 10 | .046* |
|  | R | - | - | - | - | - |
| PAG | L | - | - | - | - | - |
|  | R | 3.59 | 2 | -28 | -10 | .013* |
| LC | L | - | - | - | - | - |
|  | R | - | - | - | - | - |

*Note:* P-values are reported at peak level (family-wise-error-corrected). *Abbreviations:* RET 1 = first block of retrieval, L = left, R = right, PAG = Periaqueductal Gray, LC = Locus Coeruleus.

**Table 4.**
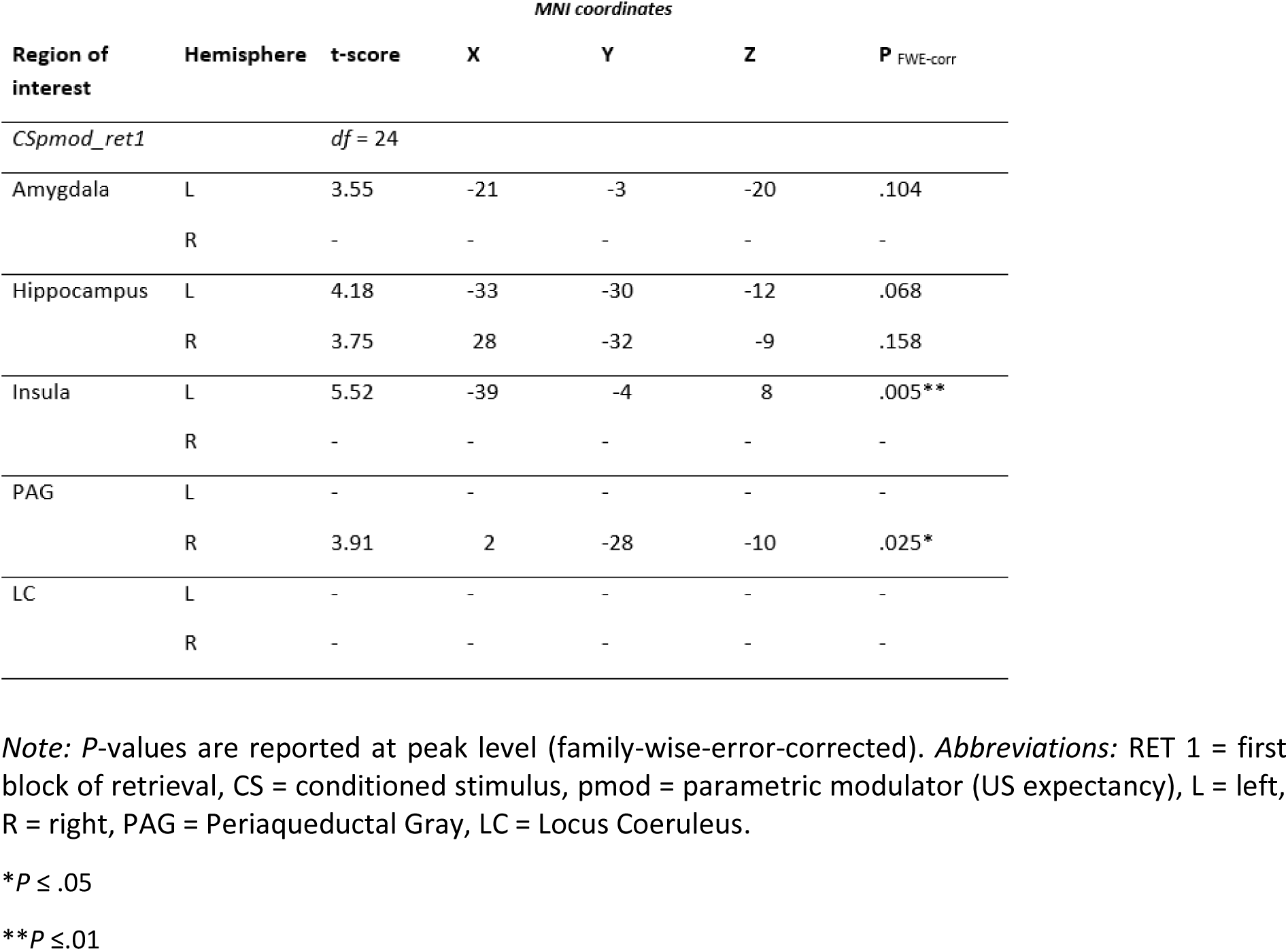
Results of the one-sample t-test (DAY 2 – RET 1), N = 25.

**Table 5.**
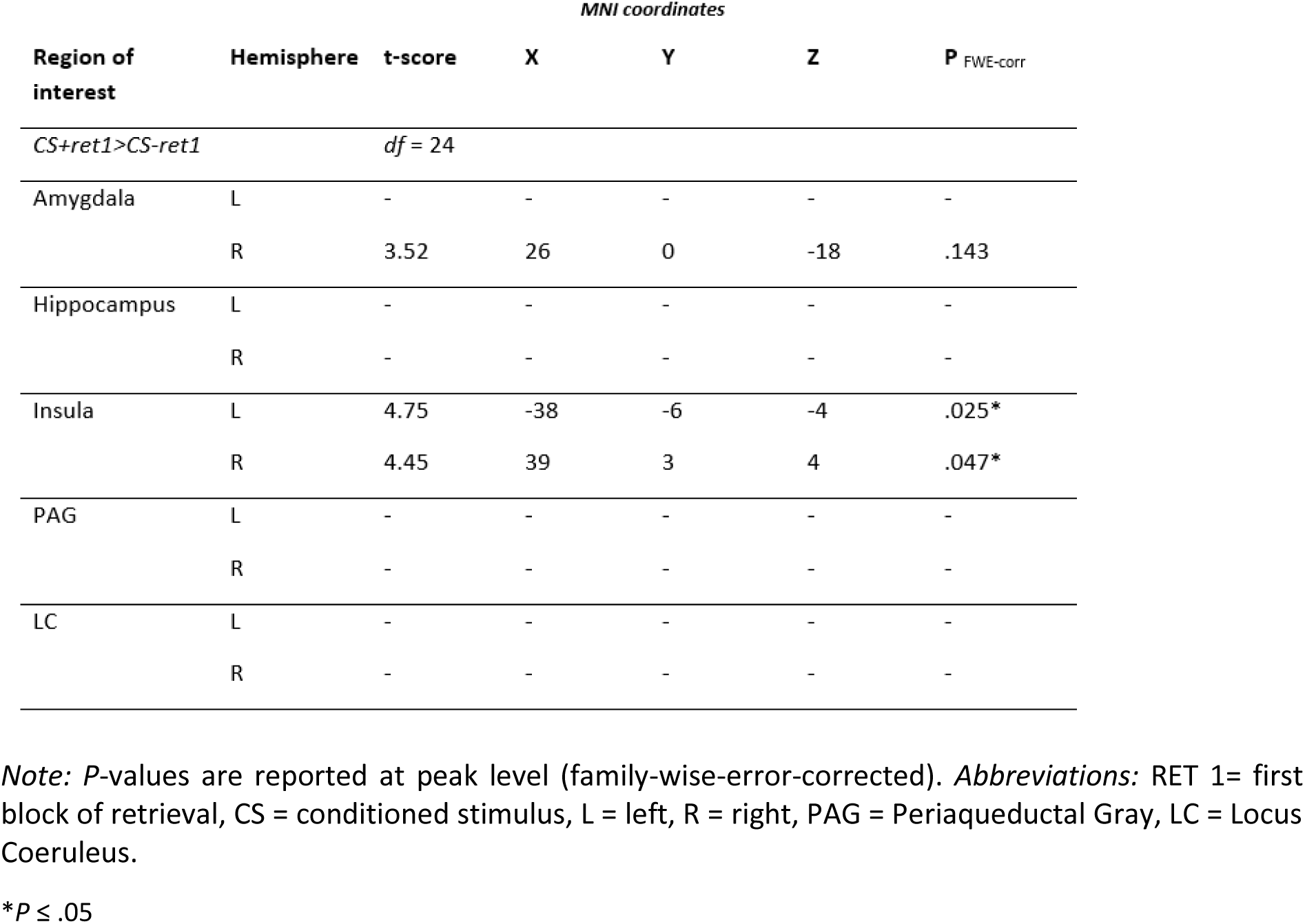
Results of physio-psychological interaction analysis (DAY 2 – RET 1), N = 25.

| Region of interest | Hemisphere | t-score | MNI coordinates |  |  | P<sub>FWE-corr</sub> |
| --- | --- | --- | --- | --- | --- | --- |
|  |  |  | X | Y | Z |  |
| CS+ret1>CS-ret1 |  | df = 24 |  |  |  |  |
| Amygdala | L | - | - | - | - | - |
|  | R | 3.52 | 26 | 0 | -18 | .143 |
| Hippocampus | L | - | - | - | - | - |
|  | R | - | - | - | - | - |
| Insula | L | 4.75 | -38 | -6 | -4 | .025* |
|  | R | 4.45 | 39 | 3 | 4 | .047* |
| PAG | L | - | - | - | - | - |
|  | R | - | - | - | - | - |
| LC | L | - | - | - | - | - |
|  | R | - | - | - | - | - |

*Note:* *P*-values are reported at peak level (family-wise-error-corrected). *Abbreviations:* RET 1= first block of retrieval, CS = conditioned stimulus, L = left, R = right, PAG = Periaqueductal Gray, LC = Locus Coeruleus.
\**P* ≤ .05

Regarding the analysis of the generalisation phase, we had to exclude four subjects from the analysis (N = 21), because of technical issues. In this phase we were interested in whether the mean US expectancy ratings also modulate the activity in our regions of interest. Results of the analysis show that the US expectancy gradient that revealed generalisation across stimuli, was reflected as hemodynamic activation in voxel in the left PAG (−2 −30 −9; *t*(21) = 4.43, *p*FWE = .012) and trend-wise in the right hippocampus (28 −30 −9, *t*(21) = 4.23, *p*FWE = .083), see *Table 6* and *Figure 12*). Additionally, we tested whether BOLD responses change with the orientation of the stimuli in the regions of interest. Particularly, activation in the bilateral insula (left insula: −34 20 0; *F*(160) = 5.16, *p*FWE = .007; right insula: 36 18 −2; *F*(160) = 5.46, *p*FWE = .004) and left PAG (−2 −34 −9; *F*(160) = 4.00, *p*FWE = .023) was associated with the identity of generalisation stimuli (see *Table 7* and *Figure 13*).

**Figure 12.**
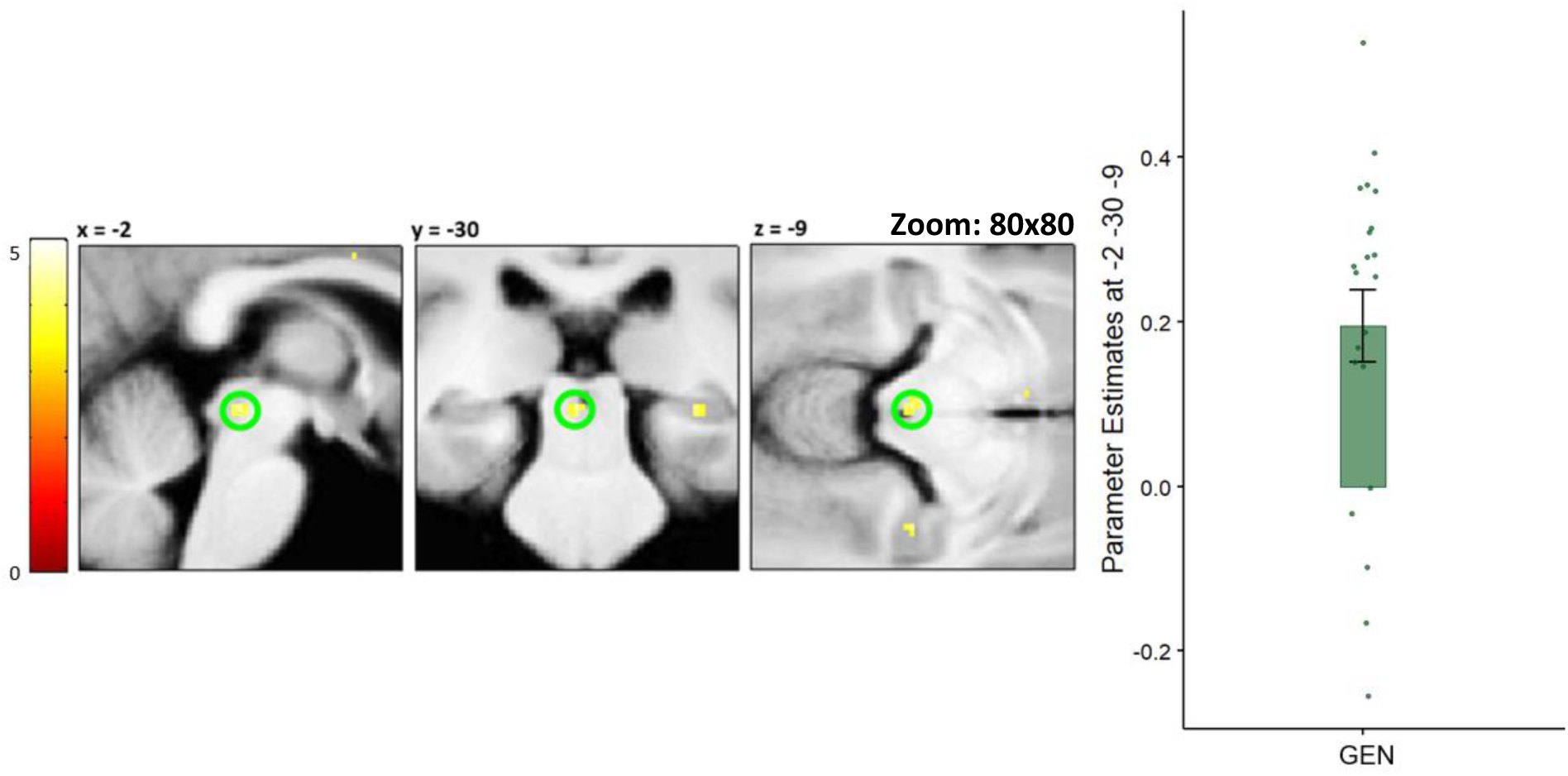
US expectancy driven modulation of BOLD activity during GEN. Results of the parametric modulation analysis during GEN show that US expectancy ratings significantly modulate BOLD activity in the PAG. *Abbreviations:* GEN = generalisation, PAG = Periaqueductal Gray.

**Figure 13.**
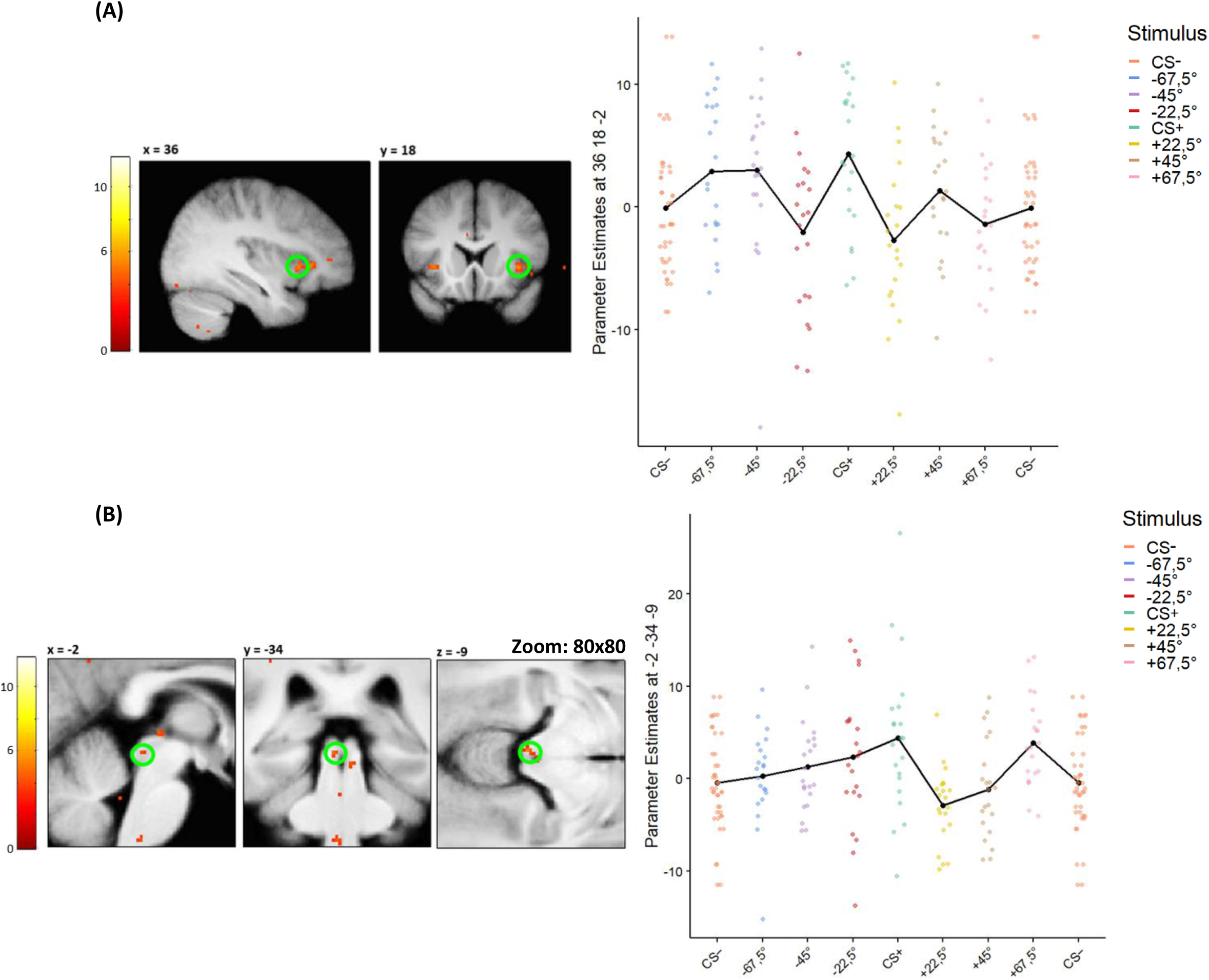
Results of full factorial analysis during GEN in (A) right insula (B) PAG. During GEN we found that the BOLD activity changes accordingly to the identity of the GEN stimuli in the **(A)** right insula and **(B)** PAG. *Abbrevations:* GEN = generalisation, PAG = Periaqueductal Gray.

**Table 6.** Results of the one-sample t-test (DAY 2 – GEN), N = 21.

| *MNI coordinates* |  |  |  |  |  |  |
| --- | --- | --- | --- | --- | --- | --- |
| Region of interest | Hemisphere | t-score | X | Y | Z | P<sub>FWE-corr</sub> |
| *genMEANPmodUSexp* |  | *df = 20* |  |  |  |  |
| Amygdala | L | - | - | - | - | - |
|  | R | - | - | - | - | - |
| Hippocampus | L | - | - | - | - | - |
|  | R | 4.23 | 28 | -30 | -9 | .083 |
| Insula | L | - | - | - | - | - |
|  | R | - | - | - | - | - |
| PAG | L | 4.43 | -2 | -30 | -9 | .012* |
|  | R | - | - | - | - | - |
| LC | L | - | - | - | - | - |
|  | R | - | - | - | - | - |

*Note:* P-values are reported at peak level (family-wise-error-corrected). *Abbreviations:* GEN = generalisation, *genMEANPmodUSexp* = parametric modulation using mean US expectancy ratings during generalisation, L = left, R = right, PAG = Periaqueductal Gray, LC = Locus Coeruleus.
\* $P \leq .05$

**Table 7.** Results of full factorial analysis (DAY 2 - GEN), N = 21.

| Region of interest | Hemisphere | F-score | MNI coordinates |  |  | P<sub>FWE-corr</sub> |
| --- | --- | --- | --- | --- | --- | --- |
|  |  |  | X | Y | Z |  |
| Effect of interest |  | df = 160 |  |  |  |  |
| Amygdala | L | - | - | - | - | - |
|  | R | - | - | - | - | - |
| Hippocampus | L | - | - | - | - | - |
|  | R | - | - | - | - | - |
| Insula | L | 5.16 | -34 | 20 | 0 | .007** |
|  | R | 5.46 | 36 | 18 | -2 | .004** |
| PAG | L | 4.00 | -2 | -34 | -9 | .023* |
|  | R | - | - | - | - | - |
| LC | L | - | - | - | - | - |
|  | R | - | - | - | - | - |

*Note:* P-values are reported at peak level (family-wise-error-corrected). *Abbreviations:* GEN = generalisation, L = left, R = right, PAG = Periaqueductal Gray, LC = Locus Coeruleus.
\* $P \leq .05$
\*\* $P \leq .01$

## DISCUSSION

Our results show convergent evidence across behaviour, physiological measures, and fMRI that participants learned, retrieved, and generalized the threat associations. During acquisition, robust CS+/CS– discrimination was reflected in subjective ratings of fear and arousal, US expectancy, as well as increased BOLD responses in bilateral insula and PAG, regions consistently implicated in interoceptive processing and defensive responding (Craig, 2009; Mobbs et al., 2009). On the following day, retrieval of the learned association was evident in both behavioural and psychophysiological outcome measures along with neural activations in the hippocampus, insula, and PAG. Functional connectivity analyses suggest that the hippocampus interacts with the insula a key hub of the salience network, to reinstate learned threat predictions (Milad & Quirk, 2012). Beyond retrieval, our data demonstrate that learned fear generalizes across similar stimuli, following graded patterns in US expectancy ratings, pupil dilation, and BOLD responses. Such generalisation gradients, which in our sample were well captured by Gaussian models, align with prior work highlighting hippocampal–salience–brainstem circuits as critical for generalisation of fear (Onat & Büchel, 2015; Dunsmoor & Paz, 2015). Together, these findings situate our paradigm within a growing body of research showing that threat associations are flexibly retrieved and generalised across modalities, engaging a coordinated cortico-limbic–brainstem network.

In our study, we find that the expectation for aversive outcomes during memory retrieval drive physiological measures of arousal (i.e. SCR and pupil-responses). Therefore, trial-wise US expectancy might provide a fine-grained parameter to explain psychophysiological modulation during emotional memory retrieval. Previous research found mixed results on modulation of SCR responses by CS contingency awareness and conscious discrimination of threat and safety (Sevenster et al., 2014); for non-findings see (Bechara et al., 1995; Esteves et al., 1994; Knight et al., 2003, 2006). However, these studies often tested the association between SCR and US expectancy ratings during threat learning phases, in which we found no modulation of psychophysiological responses during learning. Additionally, individual time-courses of US expectation further modulated neural dynamics in the insula and PAG during retrieval. Previously, it was found that the intensity of activation in the anterior insula accounted for the strength of the link between the threat-learning task and participants’ self-reported anxiety (Savage et al., 2021). Our observation of dynamic response changes in the periaqueductal gray (PAG) aligns with the findings of Roy et al., 2014, who identified this region as the core substrate for encoding aversive prediction errors. The pattern we observed, in which responses are modulated by the transition from high to low expectation during retrieval (i.e., no US present), is consistent with the axiomatic properties of a prediction error (PE) signal (since there were only non-reinforced trials during retrieval), which is largest when updating by unexpected outcomes and smallest for fully predicted ones (i.e., no updates). Our finding of insula-hippocampus coupling during retrieval may support a memory processing cycle of memory retrieval, salience detection, updating, and storage. In this model, the insula acts as a salience monitor, evaluating the accuracy of retrieved memories (Menon & Uddin, 2010). This interpretation is further supported by Wang & Yang, 2023, who identified the anterior insula as a hub for error monitoring. Their research demonstrated that the insula’s functional connectivity with the hippocampus is highly dynamic. Positive coupling helps to stabilize corrected memories, while negative coupling works to suppress incorrect ones. Therefore, our observed insula-hippocampus coupling likely reflects this adaptive mechanism, where the insula signals the hippocampus to either reinforce or revise a memory, thereby ensuring its fidelity for long-term storage. Hippocampal activity was often reported in threat-relevant encoding of contexts (Alvarez et al., 2008). In the cue-based conditioning paradigm, the confrontation with the CS+ is associated with a retrieval of a threat context that gates the cue-specific responses. The bilateral hippocampal activity observed at the beginning of retrieval may thereby also reflect the engagement of pattern separation mechanisms, i.e. separating the safety memory from the CS-from the threat associations of the CS+. As proposed by Neudert et al., 2023, this process is critical for representing similar events as distinct neural representations, which in this case would be necessary to differentiate the threat memory trace from different memory traces associated with a similar, yet different cue orientation.

Beyond retrieval, the pattern we observed during GEN suggests that the behavioural, physiological and neural expression of threat memory do not follow the categories of the stimulus identities of the CS+ and CS-, but can be described by graded responses, i.e. subjective predictions and physiological responses extend beyond the originally predictive cue. We found a Gaussian-like decay of US expectancy and pupil responses, which may support the suggestion that the brain maintains a continuous representation of stimulus similarity to estimate threat probability (Dunsmoor & Paz, 2015; Shepard, 1987). However, importantly to note here is that the mechanism by which generalisation occurs may not be driven by the inability to perceptually discriminate between stimuli, but a dynamic process in which categorical conceptualisations and prior associations shape response gradients (Dunsmoor & Murphy, 2015; Onat & Büchel, 2015). This also aligns with our results that US expectancy ratings modulate BOLD activity in the PAG and hippocampus. The PAG output has been linked to defensive responses (Mobbs et al., 2007) and its correlation with subjective expectancy suggests that even in absence of a US, participants’ learned model of threat probability continues to shape the current responses in the defensive circuitry. This may be supported by the trend-wise hippocampal modulation by US expectancy, which may encode and retrieve a “similarity space” for stimuli (Cooper & Ritchey, 2020; Shepard, 1987). We find insula activation during the categorisation of stimulus identity, which may be supported by the results of (Onat & Büchel, 2015), in which they found that the more the similarity to the CS+ decreased, the more the activation of the insula decreased. This is in line with the literature that the insula activity represents CS similarity (Dunsmoor & Paz, 2015; Fullana et al., 2016; Lange et al., 2017; Lissek et al., 2014). By showing convergent gradients in US expectancy and pupil responses and linking them to PAG and insula signals, our findings extend prior work that often relied on single modalities and suggest multimodal outcome measures could be valuable as sensitive markers of individual generalisation profiles.

Although the present multimodal approach provides a rich picture of fear learning, retrieval and generalisation, several methodological limitations need to be considered. Firstly, we did not find robust amygdala effects despite its well established role in conditioned fear (Sehlmeyer et al., 2009). Because the medial temporal lobe is vulnerable to susceptibility artifacts, acquiring reliable data can be challenging. This may lead to compromised image integrity and a resulting loss of functional BOLD signal in this region (Bellgowan et al., 2006; Morawetz et al., 2008; Sehlmeyer et al., 2009). Secondly, SCR, although showing generalisation, were poorly captured by Gaussian modelling. This likely reflects that SCR is an unspecific and slowly evolving autonomic outcome measure, which rather depends on cognitive awareness and contingency evaluation than continuous perceptual similarity (Bach et al., 2010). However, the set-up of our experiment was not tailored to capture these fine-graded responses. SCR thus may act as a bottleneck: information must pass through cognitive appraisal (making it “slow” and “cognitive”), but the output system itself lacks the precision of the original neural and perceptual representations, which results in a broad, non-selective measure of arousal rather than a sharp indicator of fine perceptual gradients (Sevenster et al., 2014). Thirdly, although our paradigm models generalisation along a perceptual continuum, emerging work suggests that overgeneralisation in anxiety is not simply perceptual blurring but involves integration failures of prior knowledge and conceptual boundaries (Cha, Carlson, et al., 2014; Cha, Greenberg, et al., 2014). Extending our approach to manipulations of conceptual category boundaries would allow testing whether the hippocampal-salience-brainstem network we identified shows is also involved in altered integration and therefore, overgeneralisation. Finally, our sample size was modest and exclusions for physiological noise further reduced power for SCR and pupil analyses. Future studies with larger and more diverse samples could improve reliability and allow approaches to detect individual differences (e.g. linking gradient width to trait anxiety).

